# The ancestral ESCRT protein TOM1L2 selects ubiquitinated cargoes for retrieval from cilia

**DOI:** 10.1101/2022.09.23.509287

**Authors:** Swapnil Rohidas Shinde, David U. Mick, Erika Aoki, Rachel B. Rodrigues, Steven P. Gygi, Maxence V. Nachury

**Affiliations:** Department of Ophthalmology, University of California San Francisco, CA 94143, USA; Center of Human and Molecular Biology and Center for Molecular Signaling, Department of Medical Biochemistry and Molecular Biology, Saarland University School of Medicine, Homburg, Germany; Department of Cell Biology, Harvard Medical School, Boston, MA 02115, USA

## Abstract

Many G protein-coupled receptors (GPCRs) reside within cilia of mammalian cells and must undergo regulated exit from cilia for the appropriate transduction of signals such as Hedgehog morphogens. Lysine 63-linked ubiquitin (K63Ub) chains mark GPCRs for regulated removal from cilia, but the molecular basis of K63Ub recognition inside cilia remains elusive. Here we show that the BBSome –the trafficking complex in charge of retrieving GPCRs from cilia– engages the ancestral endosomal sorting factor TOM1L2 (Target of Myb1-Like 2) to recognize UbK63 chains within cilia. TOM1L2 directly binds to UbK63 chains and to the BBSome and targeted disruption of the TOM1L2/BBSome interaction results in the accumulation of TOM1L2, ubiquitin and the GPCRs SSTR3, Smoothened and GPR161 inside cilia. Strikingly, the single cell alga *Chlamydomonas* also requires its TOM1L2 orthologue to clear ubiquitinated proteins from cilia. We conclude that TOM1L2 broadly enables the retrieval of UbK63-tagged proteins by the ciliary trafficking machinery.

## INTRODUCTION

Primary cilia transduce sensory, developmental, and homeostatic signals by dynamically concentrating signaling receptors together with their downstream signaling machinery (Anvarian et al., 2019; Carter and Blacque, 2019; Nachury and Mick, 2019). For instance, addition of the Hedgehog morphogen to cells triggers the enrichment of the G protein-coupled receptor (GPCR) Smoothened (SMO) in cilia and the departure of the GPCR GPR161 from cilia (Gigante and Caspary, 2020). This redistribution of ciliary GPCRs modifies the levels of ciliary second messengers, ultimately altering the activity of transcription factors that shape the Hedgehog response. The paradigm established by Hedgehog signaling is likely to have broad significance as over 30 GPCRs have been found to reside in cilia, and in most studied instances ciliary GPCRs undergo regulated trafficking in and out of cilia (Green and Mykytyn, 2014; Hilgendorf et al., 2016). Because SMO constitutively enters and exits cilia in unstimulated cells, and because pathway activation reduces the ciliary exit rate of SMO (Milenkovic et al., 2015; Ye et al., 2018), regulated exit is responsible –at least in part– for the dynamic ciliary enrichment of SMO.

We know that GPCR exit from cilia is carried out by the BBSome, a complex of 8 Bardet-Biedl Syndrome (BBS) proteins, together with the intraflagellar transport (IFT) complexes A and B and the microtubule motor dynein-2 but regulated cargo selection by the ciliary exit machinery had remained poorly understood until recently (Nachury, 2018; Wingfield et al., 2018). The attachment of the small polypeptide ubiquitin onto proteins modifies their fate, often towards degradative destinies, and the specific linkage used to elongate ubiquitin chains drives the specific biological outcome. Lysine 63-linked ubiquitin (UbK63) chains mark membrane proteins for sorting to the lysosome and we and others recently showed that UbK63 chains mark GPCRs for removal from cilia (Desai et al., 2020; Shinde et al., 2020). Activation of the somatostatin receptor 3 (SSTR3) leads to its arrestin-dependent ubiquitination and subsequent exit (Shinde et al., 2020) while constitutive ubiquitination of SMO by the ligase WWP1 keeps ciliary SMO levels low under basal conditions (Lv et al., 2021). For both SSTR3 and SMO as well as for GPR161, targeted cleavage of UbK63 chains inside cilia by the UbK63 deubiquitinase AMSH blocks ciliary exit (Desai et al., 2020; Shinde et al., 2020).

The biological importance of UbK63 chains in ciliary exit now poses the question of how UbK63 chains are recognized by the ciliary exit machinery. One possibility is that ciliary trafficking complexes directly recognize ubiquitin. The BBSome was recovered over immobilized ubiquitin from trypanosome extracts (Langousis et al., 2016) and IFT139 was found to bind to ubiquitinated tubulin (Wang et al., 2019). However, no direct binding to Ub has been established for the BBSome or IFT-A nor has selectivity for K63-linked chains been tested. Alternatively, candidate ciliary UbK63 readers may be found amongst the well-established UbK63 readers that recognize and sort membrane proteins to the lysosome and autophagosome (Grumati and Dikic, 2018; Piper et al., 2014). The endosomal sorting complexes required for transport (ESCRT) comprise successively acting protein complexes that sort membrane proteins marked with UbK63 chains from the limiting membrane of endosomes into intralumenal vesicles destined for lysosomal degradation (Schöneberg et al., 2017; Vietri et al., 2020). In nematodes, mutations in either the canonical ESCRT-0 complex HRS/STAM or in the BBSome cause the accumulation of Ub-fusion membrane proteins inside cilia (Hu et al., 2007; Xu et al., 2015), suggesting that HRS/STAM may participate in ciliary sorting of ubiquitinated proteins by the BBSome. Alternatively, a coupling between ciliary exit and degradation may underlie the requirement for ESCRT-0 in ciliary exit.

Here, we conduct biochemical assays and find that neither the BBSome nor the IFT-A machinery directly recognize UbK63 chains. Instead, proteomics profiling, focused screens and biochemical mapping reveal that the ancestral ESCRT-0 protein TOM1L2 (target of Myb1-like 2) is the adaptor that bridges the BBSome to its ubiquitinated cargoes.

## RESULTS

### The BBSome needs an adaptor to recognize UbK63 chains

As prior studies hinted at possible interactions between ubiquitin and either the BBSome or IFT-A (Langousis et al., 2016; Wang et al., 2019), we directly tested whether the BBSome and the IFT complexes can associate with UbK63 chains. Because Ub chains remain attached to HECT-type ubiquitin ligases and because HECT ligases build chains with a high degree of specificity, one can grow Ub chains with defined linkages onto the immobilized catalytic domain of HECT ligases to conduct capture assays (Nathan et al., 2013). Sepharose-bound GST-NEDD4 was incubated with the required ubiquitination machinery to assemble UbK63 chains, proteins were captured onto GST-NEDD4-UbK63 resin and UbK63-binding proteins were specifically eluted by cleaving the Ub chains with the deubiquitinase Usp2 (**Fig. 1A**). To control for non-specific binding to GST-NEDD4, Ub was omitted from the ubiquitination reaction. Retinal extracts were used as starting material because of the high abundance of IFT complexes and BBSome. Captures from bovine retinal extracts or IMCD3 cell lysates recovered the known UbK63 readers Myosin VI, TOM1 (target of Myb1) and TOM1L2 (Nathan et al., 2013; Penengo et al., 2006; Yamakami et al., 2003) (**Fig. 1B and S1A**) and bacterially expressed TOM1L2 bound to UbK63 chains in this assay (**Fig. S1B**). In contrast, no specific binding of IFT-A or IFT-B complexes to UbK63 chains was detected (**Fig. 1B**). Similarly, no binding of the BBSome to UbK63 chains was detected from either retinal extract (**Fig. 1C**) or highly purified and concentrated BBSome preparations (**Fig. 1D**). To further test whether the BBSome may recognize UbK63 chains, we established stable cell lines expressing GFP-tagged BBSome in IMCD3 cells. Again, no binding of the BBSome to UbK63 chains was detected even though the known UbK63 reader TOM1L2 did bind in this assay (**Fig. 1E**). Together, these binding studies fail to support models where the core ciliary trafficking complexes directly recognizes the UbK63 chains attached to exiting signaling receptors. Instead, the ciliary trafficking machinery must engage adaptors to select UbK63-marked cargoes.

**Figure 1.**
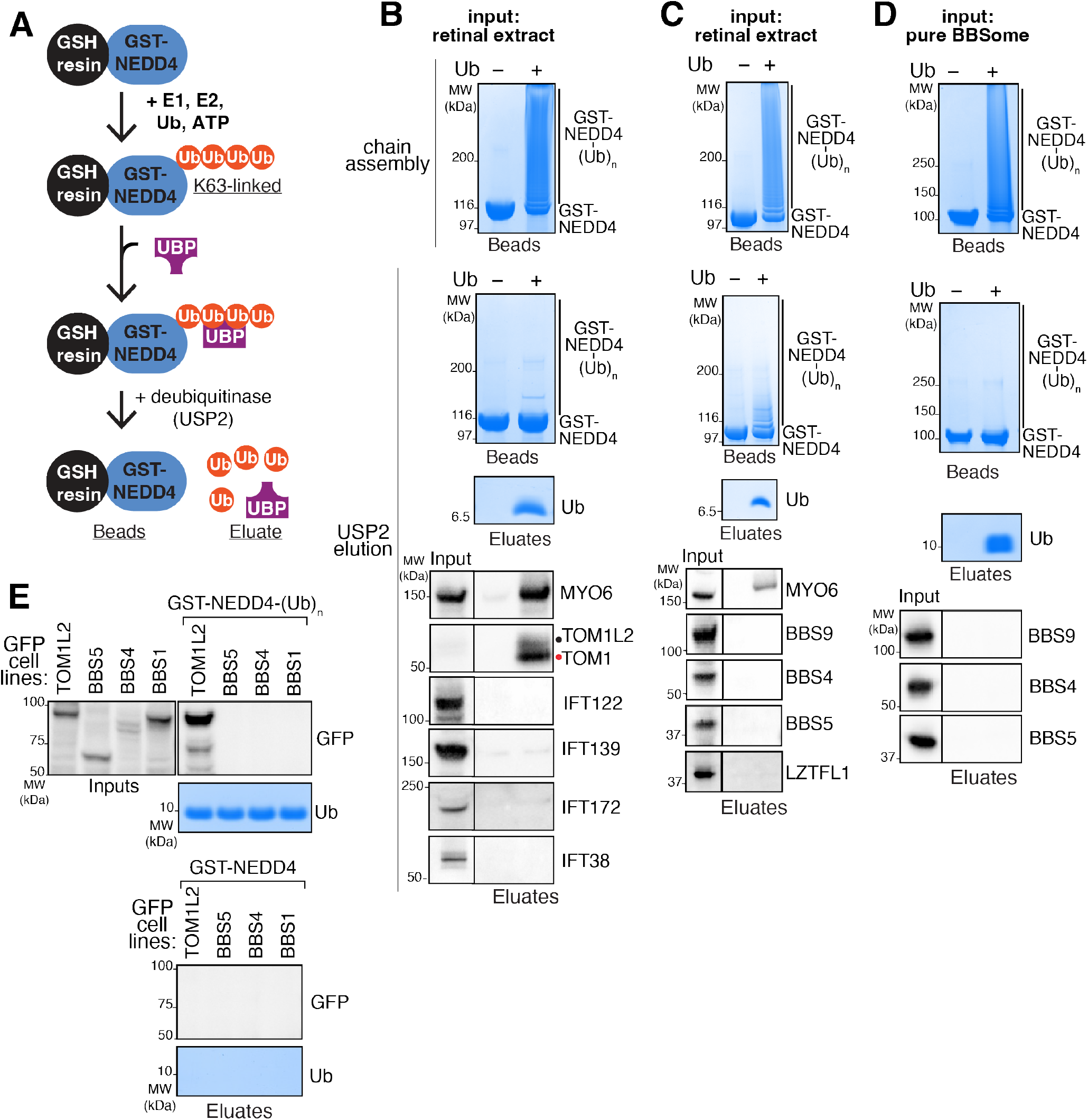
Absence of detectable binding of the BBSome and IFT complexes to UbK63 chains. **A.** Diagram of the biochemical strategy. Recombinant GST-NEDD4 bound to glutathione (GSH) resin is incubated with E1 (UBA1), E2 (UBCH5), ATP, and ubiquitin to grow UbK63 polyubiquitin chains onto NEDD4. UbK63 chains grown on immobilized NEDD4 capture ubiquitin-binding proteins (UBP) from cell or tissue extracts, and proteins bound to UbK63 chains are specifically eluted by cleaving ubiquitin chains with the deubiquitinase USP2. **B-D**. Top panel: Coomassie-stained gel of bead-bound GST-NEDD4 after completion of autoubiquitination reaction. Ub was omitted from the ubiquitination reaction for the control. Middle and lower panels show the products of the treatment with USP2. Middle panel: Coomassie-stained gel of bead-bound GST-NEDD4 ± (Ub)_n_ shows that Ub chains are fully digested by USP2. Lower panel: Coomassie-stained gel of material released from beads by USP2 treatment. Ubiquitin is only present when chains had been assembled on NEDD4. Bottom panels: western blots of known UBP and ciliary trafficking components. 0.1% input and 42% of cleavage eluates were loaded for the western analyses. In **B** and **C**, bovine retinal extracts were applied onto GST-NEDD4 ± (Ub)_n_ beads. Myosin VI (MYO6) and TOM1/TOM1L2 are known UbK63 readers and efficient binding is detected. No binding to UbK63 is detected for the IFT-A complex subunits IFT122 and IFT139, the IFT-B complex subunits IFT172 and IFT38, the BBSome subunits BBS9, BBS4 and BBS5, or the BBSome-associated protein LZTFL1. In **D**, BBSome purified from bovine retina was applied onto GST-NEDD4 ± (Ub)_n_ beads. **E.** IMCD3 cells stably expressing GFP-tagged TOM1L2, BBS5, BBS4, or BBS1 were lysed, and extracts were passed over GST-NEDD4 ± (Ub)_n_ beads. Lower panels: Coomassie-stained gel showing Ub release by USP2. Upper panels: Immunoblotting for GFP reveals specific binding of TOM1L2 to UbK63 chains, but not of the BBSome subunits BBS1, BBS4, or BBS5. GST-NEDD4 without Ub in the chain building reaction served as a control.

### The UbK63 reader TOM1L2 is required for BBSome-mediated retrieval of ciliary GPCRs

A UbK63 adaptor that bridges ubiquitylated cargoes to the retrieval machinery should accumulate in cilia together with ubiquitinated cargoes when retrieval is compromised. BBSome-mediated retrieval requires the small GTPase ARL6/BBS3, and we conducted proteomic profiling of *Arl6*^−/−^ vs. WT cilia in triplicate using the cilia-APEX platform (Mick et al., 2015) to identify candidate ciliary UbK63 retrieval adaptors (**Fig. 2A**). Among 20 proteins that accumulate at least 2-fold and with a *p*-value of 0.05 or less, four known UbK63 readers were found, TOM1, TOM1L2, TOLLIP and HRS. TOM1 family members and HRS are both ESCRT-0 proteins that function in the earliest step of ubiquitylated cargo recognition at the surface of endosomes (Piper et al., 2014; Roach et al., 2021), and TOLLIP is a known partner of TOM1 (Yamakami et al., 2003). HRS’s partner STAM2 was also found enriched in the ciliary proteome of *Arl6*^−/−^ cells, albeit with a weaker significance value. Staining *Arl6*^−/−^ cells with an antibody that recognizes TOM1 and TOM1L2 revealed a strong enrichment of TOM1/TOM1L2 in cilia when retrieval is compromised (**Fig. 2B-C**). Similarly, TOLLIP accumulates in cilia of ARL6-depleted cells (**Fig. S1C-D**). Furthermore, IFT27/BBS19 is another GTPase essential for BBSome function and proteomics profiling of *Ift27*^−/−^ cilia had previously detected an enrichment of TOM1, TOM1L2, TOLLIP as well as the TOM1/TOM1L2 partner and known UbK63 reader MYO6 (Mick et al., 2015). We previously confirmed that MYO6 accumulates in *Bbs19*^−/−^ cilia and found that it participates in ciliary GPCR shedding into extracellular vesicles, a process known as ectocytosis (Nager et al., 2017). Together, these findings point to ESCRT-0 proteins and their partners as candidates for the UbK63 retrieval adaptors.

**Figure 2.**
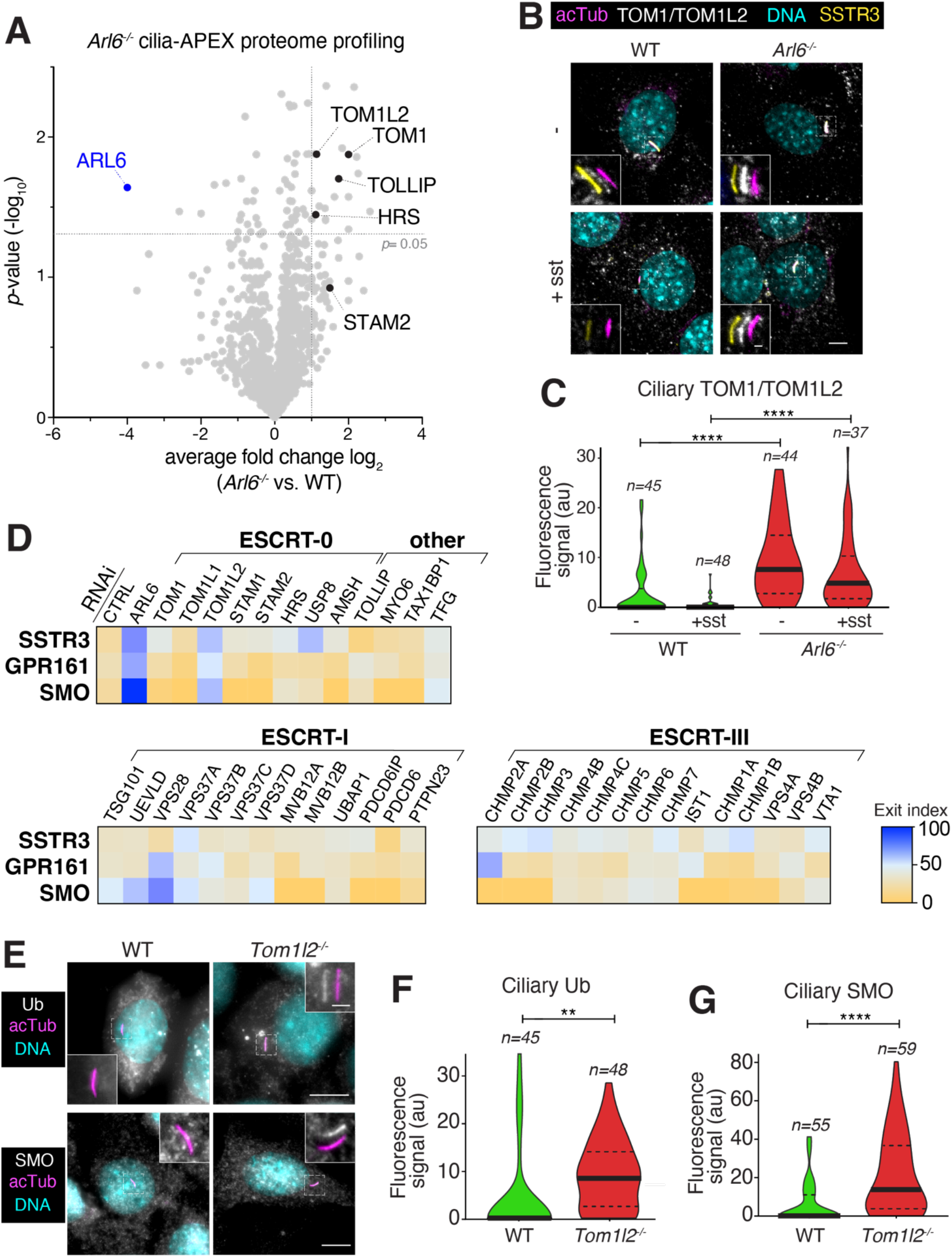
Identification of TOM1L2 as a ciliary retrieval factor by proteomics profiling and functional screening. **A.** Wildtype (WT) and *Ar16*^−/−^IMCD3 cells were subjected to Cilia-APEX proteomics. Hits are presented in a volcano plot of statistical significance versus protein enrichment. Known UbK63 readers enriched in the *Ar16*^−/−^ cilia proteome are shown as black dots. The complete data set is found in Table S1. **B.** WT or *Ar16*^−/−^ IMCD3 cells stably expressing SSTR3 fused to mNeonGreen (NG) were treated with or without somatostatin-14 (sst) for 2 h, then fixed and stained for acetylated tubulin (acTub, magenta) and TOM1/TOM1L2 (white). SSTR3^NG^ was imaged through the intrinsic fluorescence of NG (Yellow). In these and every subsequent insets, channels are shifted to facilitate visualization of ciliary signals. Scale bar: 5μm (main panel), 1μm (inset). **C.** The ciliary fluorescence intensity of the TOM1/TOM1L2 channel was measured in each condition, and the data are represented as violin plots. In this and every subsequent violin plot, the dotted lines indicate the first and third quartiles and the thick bar is the median. Asterisks indicate statistical significance values calculated by one-way ANOVA followed by Tukey’s post hoc test. ****, *p* ≤ 0.0001. **D.** IMCD3 cells stably expressing SSTR3^NG^ or GPR161^NG^ were transfected with indicated siRNAs for 48h. IMCD3-[SSTR3^NG^] cells were treated with or without somatostatin-14 (sst) for 6 h, fixed, and stained for acetylated tubulin. IMCD3-[GPR161^NG^] cells were treated with or without SAG for 3 h, fixed and stained for acetylated tubulin and SMO. SSTR3^NG^ and GPR161^NG^ were imaged through the intrinsic fluorescence of NG. The fluorescence intensities of SSTR3^NG^, GPR161^NG^, and SMO in the cilium were measured in each condition. Median fluorescence intensities with sst (SSTR3^NG^) or SAG (GPR161^NG^) treatment were normalized to untreated conditions and are represented as exit index on the heat map (see Methods for details). For SMO, median fluorescence intensities in untreated conditions are represented as exit index on the heat map. The depletion efficiencies of representative proteins are shown in **Fig. S2A**. The measurement of ciliary fluorescence for each GPCR are shown in **Fig. S3-5**. **E.** Ciliated WT or *Tom1l2*^−/−^ cells were fixed and stained for acetylated tubulin (acTub, magenta) and Ub (white) or SMO (white). For the Ub stain, cells were incubated with SAG for 2h prior to fixation to induce GPR161 exit. Scale bar: 5μm (main panel), 1μm (inset). **F.** The fluorescence intensity of the ubiquitin channel in the cilium was measured in each condition, and the data are represented as violin plots. Asterisks indicate unpaired *t*-test significance value. **, *p* ≤ 0.01. *n* = 45-48 cilia. **G.** The fluorescence intensities of ciliary SMO are represented as violin plots. Asterisks indicate unpaired *t*-test significance value. ****, *p* ≤ 0.0001. *n* = 55-59 cilia.

In addition, we noticed that several ESCRT proteins were identified in the cilia-APEX2 proteome (TSG101, CHMP4B, VPS37B/C) (May et al., 2021) and in the proteomic profiling of *Bbs17*^−/−^ photoreceptor cilia (VPS4/VTA1, VPS28, CHMP5) (Datta et al., 2015). As UbK63 recognition is broadly distributed among the ESCRT complexes, we set out to test the role of all ESCRT-0, ESCRT-I and ESCRT-III in GPCR retrieval from cilia. Additional UbK63 readers identified in previous ciliary proteomics studies were also included (e.g. MYO6, TAX1BP1). To avoid identifying components that may only function in one signaling modality or regulate a specific GPCR, we assessed the exit of three GPCRs with distinct regulatory inputs but that all require UbK63 chains for their exit from cilia (Desai et al., 2020; Shinde et al., 2020). SMO undergoes constitutive exit from cilia under basal signaling conditions, GPR161 exits cilia upon activation of the Hedgehog pathway, and SSTR3 represents the prototype of a ciliary GPCR that undergoes exit when stimulated by its agonist somatostatin. To focus the screen on retrieval, we blocked the alternative exit path of ectocytosis during the exit assays with low doses of the actin poison cytochalasin D (Nager et al., 2017; Phua et al., 2017).

Screening of 40 ESCRT and UbK63-related candidates for exit of GPR161, SSTR3 and SMO by siRNA revealed TOM1L2 as the single common hit (**Fig. 2D and S2A**). The only other hit affecting more than one GPCR was the ESCRT-I subunit VPS28 whose depletion interfered with GPR161 and SMO exit but not with SSTR3 exit. We noted that depletion of VPS28 led to a drastic accumulation of ubiquitin and of TOM1/TOM1L2 on endosomal structures, suggesting that VPS28 depletion may indirectly affect retrieval by trapping TOM1L2 on endomembrane compartments (**Fig. S2C**). Furthermore, we found no evidence of VPS28 presence in cilia by immunostaining (**Fig. S2B**) or by Cilia-APEX (May et al., 2021; Mick et al., 2015) and **Table S1**). We thus consider it unlikely that VPS28 directly participates in ciliary retrieval. We note that MYO6, a protein previously implicated in ectocytosis of ciliary GPCRs, did not score in the retrieval screens despite being efficiently depleted by siRNA (**Fig. S2A**). Furthermore, we note that depletion of HRS or STAM fail to block the regulated ciliary exit of GPCRs indicating that blockage of endolysosomal sorting is not sufficient to block exit from cilia through a hypothetical coupling between the two processes.

To validate the function of TOM1L2 in ciliary retrieval of ubiquitinated GPCRs, we generated a knockout cell line via CRISPR/Cas9 (**Fig. S1E**) and stained for Ub and SMO (**Fig. 2E-G**). Both SMO and Ub accumulated in cilia of *Tom1l2*^−/−^ cells compared to WT cells, where their levels were largely undetectable. Furthermore, signal-dependent exit of GPR161 was severely compromised in *Tom1l2*^−/−^ cells (**see Fig. 7A**). Together, these results support a role for TOM1L2 in the regulated exit of ciliary GPCRs.

**Figure 3.**
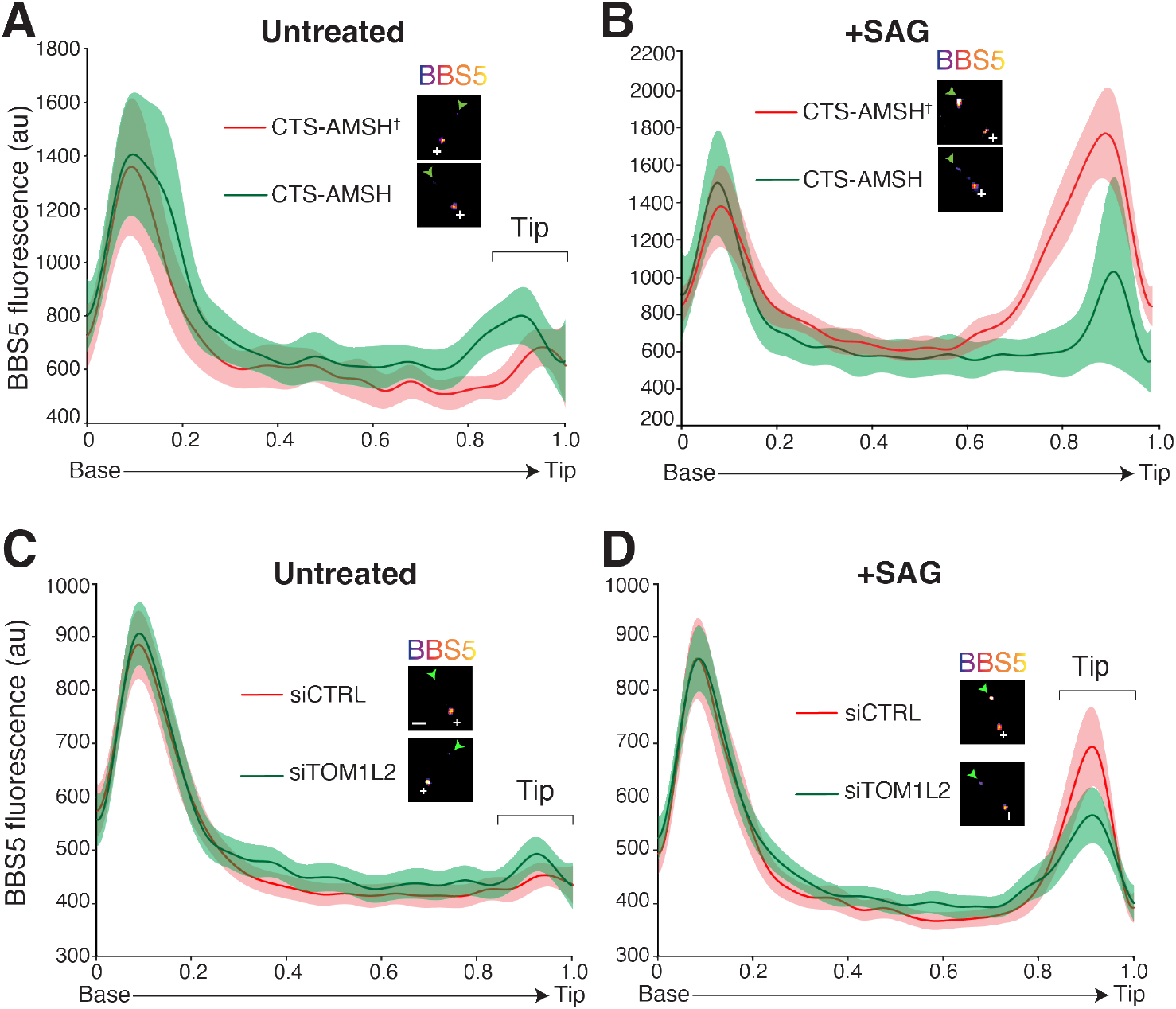
TOM1L2 functions between the BBSome and ubiquitin recognition. Line scans of ^NG3^BBS5 fluorescence intensities along cilia and corresponding micrographs. Representative images of cilia are shown in insets. Scale bar: 1μm. ^NG3^BBS5 is in fire scale, a white cross marks the location of the base, and an arrowhead marks the tip of the cilia. In all line scans, the line marks the mean intensity along length-normalized cilia and the shaded area shows the 95% confidence interval. **A-B.** Control- and UbK63-depleted cilia. IMCD3-[pEF1α-^NG3^BBS5] were transfected with plasmids expressing the ciliary targeting signal (CTS) of NPHP3 fused to mScarlet and either the catalytic domain of the K63-specific deubiquitinase AMSH, or the catalytically inactive E280A variant (AMSH†). Cells were serum starved and then left untreated (**A**) or treated with SAG (**B**) for 40 min before fixation and TIRF imaging of NeonGreen fluorescence. See **Fig. S6A** for additional images. *n* = 9-11 cilia. **C-D.** Control- and TOM1L2-depleted cells. IMCD3-[pEF1α-^NG3^BBS5] were transfected with non-targeting control siRNAs (siCTRL) or TOM1L2 siRNAs (siTOM1L2), serum-starved. and either left untreated (**C**) or treated with SAG (**D**) for 40 min. Cells were then fixed and NeonGreen fluorescence was imaged by TIRF. See **Fig. S6C** for additional images. *n* = 30-45 cilia.

**Figure 4.**
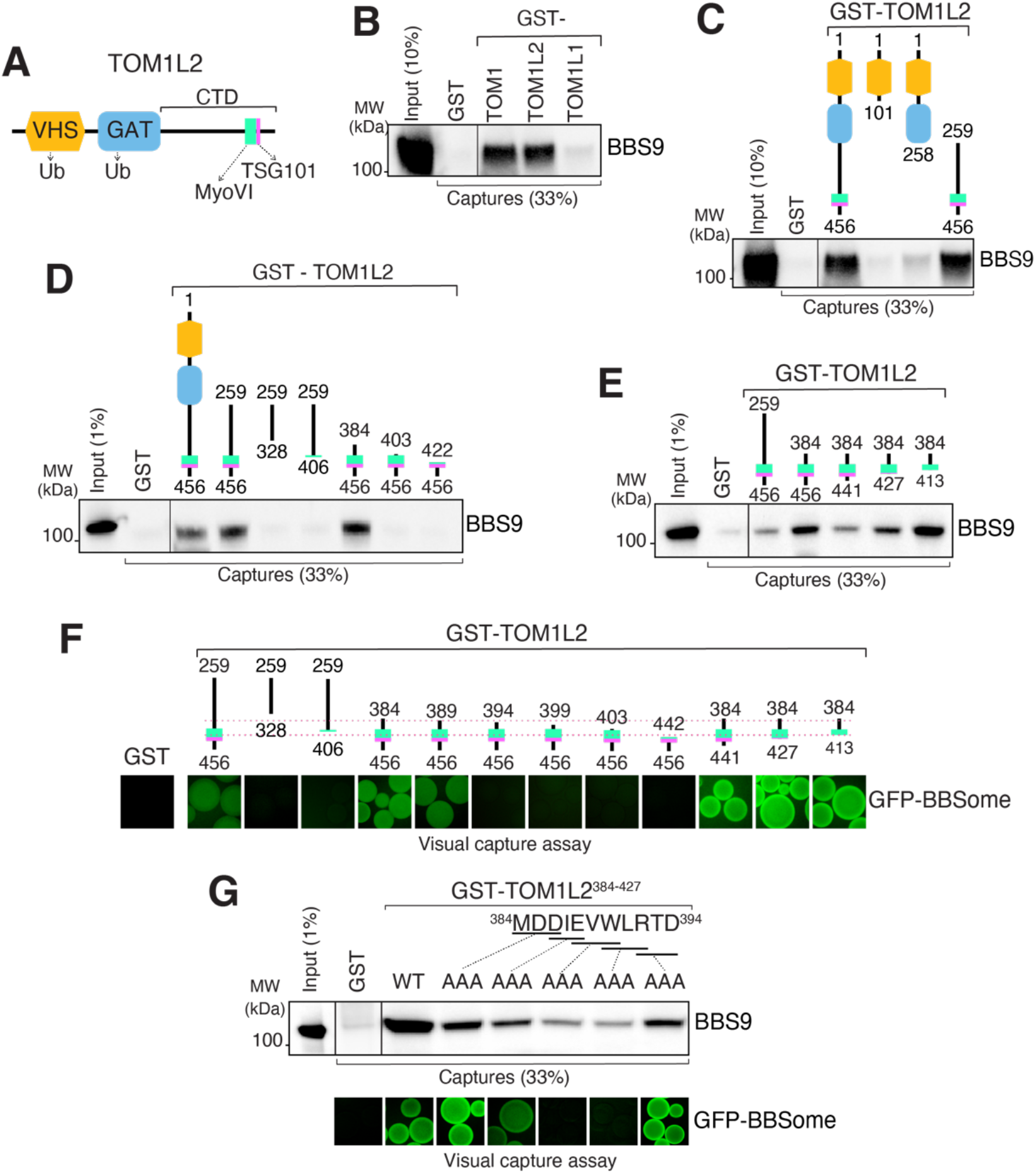
Mapping of the BBSome-binding determinant on TOM1L2. **A.** Diagram of the domain organization of TOM1L2. **B.** GST-TOM1, GST-TOM1L2 and GST-TOM1L1 were immobilized on glutathione sepharose and beads incubated with BBSome purified from the bovine retina. Bound material was eluted in SDS sample buffer and the BBSome was detected by immunoblotting for BBS9 (and BBS4 and BBS5, see **Fig. S7A**). The purity of the GST fusion proteins is shown in the Ponceau stains found in **Fig. S7A**. **C.** Capture assays of pure BBSome were conducted with truncations of TOM1L2 fused to GST. See **Fig. S7B** for additional immunoblots and Ponceau stain. **D-E.** Capture assays with truncations of TOM1L2 find that aa 384-413 are necessary and sufficient for binding to the BBSome. **F.** Visual capture assays were conducted with truncations of TOM1L2 fused to GST and immobilized onto glutathione sepharose and extracts from HEK cells overexpressing all eight BBSome subunits fused to GFP. **G.** TOM1L2 triple alanine mutants were fused to GST and immobilized onto glutathione sepharose and used for conventional capture assays with BBSome purified from bovine retinas (upper panel) or for visual capture assays with extracts from HEK cells overexpressing all eight BBSome subunits fused to GFP (lower panel).

**Figure 5.**
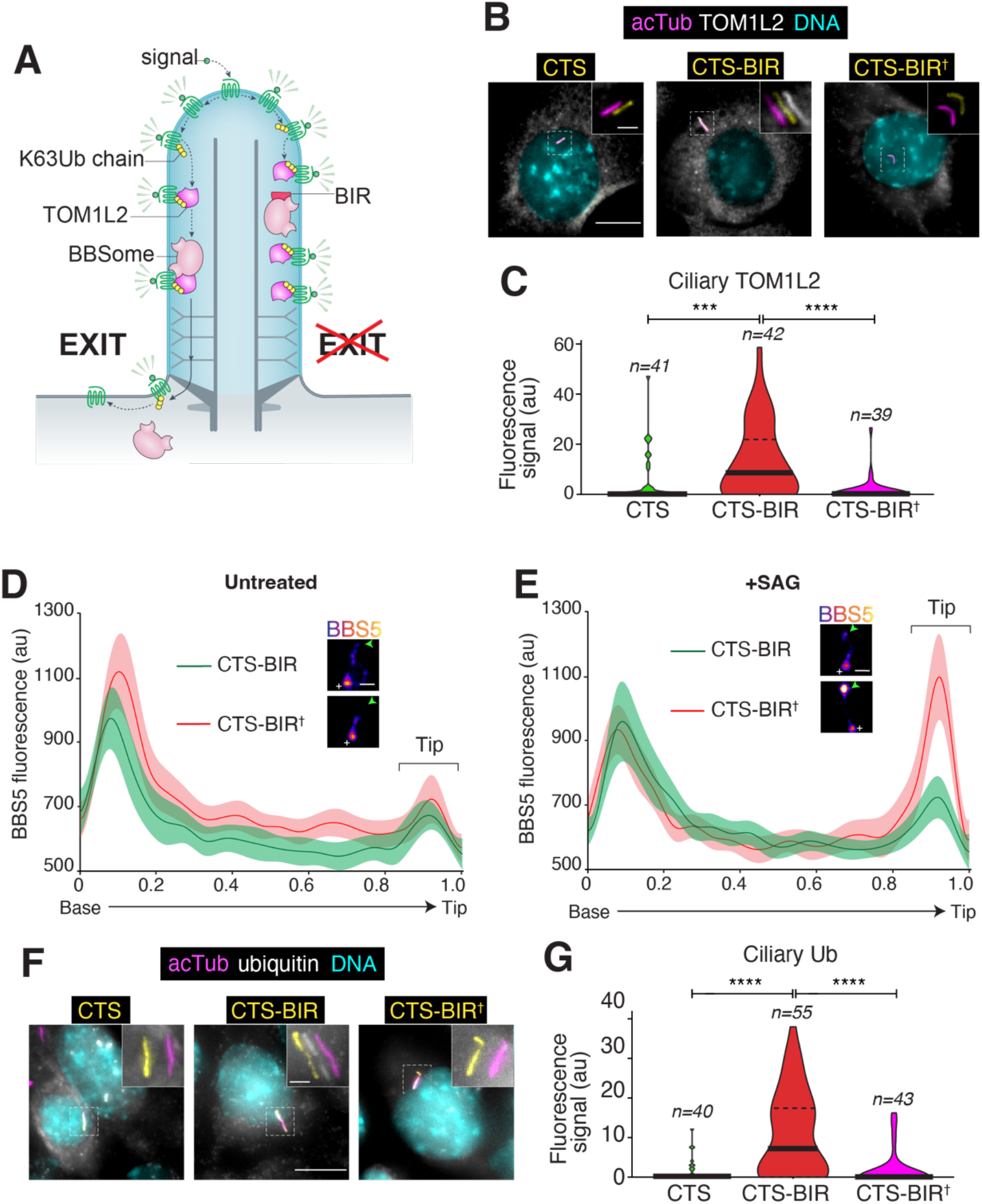
Targeted disruption of the BBSome-TOM1L2 interaction blocks the exit of ubiquitinated proteins and of TOM1L2 from cilia. **A.** Diagram of the working model and the experimental strategy. **B.** IMCD3-[pEF1α-TOM1L2^FLAG3^] cells were transfected with plasmids expressing the ciliary targeting signal (CTS) of NPHP3 fused to GFP, or CTS fused to GFP and the BBSome binding motif of TOM1L2 (BIR), or to GFP and the ^390^WLR^392^/AAA mutant of the BIR defective in BBSome binding (BIR†). Cells were serum-starved 24h later, fixed after another 24h, and stained for acetylated tubulin (acTub; magenta), FLAG (TOM1L2, white), and DNA (cyan). The CTS fusions were visualized through the intrinsic fluorescence of GFP (yellow). Scale bars, 5 μm (main panel), 1 μm (inset). **C.** The fluorescence intensities of ciliary TOM1L2^FLAG3^ are represented as violin plots. *n* = 39-42 cilia. Asterisks indicate significance values for Tukey’s multiple comparison test. ****, *p* ≤0.0001; ***, *p* ≤ 0.001. **D-E.** Line scans of ^NG3^BBS5 fluorescence intensities along cilia. IMCD3-[pEF1α-^NG3^BBS5] cells transfected with the indicated plasmids were either left untreated (**D**) or treated with SAG (**E**) for 40 min, fixed, and imaged. Representative images of cilia are shown in insets (see also **Fig S6B**). Scale bar: 1μm. ^NG3^BBS5 is in fire scale, white crosses mark the location of the basal body, and an arrowhead marks the tip of the cilia. *n* = 14-20 cilia. **F.** IMCD3 cells transfected with the indicated plasmids were treated with SAG for 2h, fixed, and stained for acetylated tubulin (acTub; magenta), ubiquitin (Ub, white), and DNA (cyan). The CTS fusions were visualized through the intrinsic fluorescence of GFP (yellow). Scale bars, 5 μm (main panel), 1 μm (inset). **G.** The fluorescence intensities of ubiquitin in cilia are represented as violin plots. *n* = 40-55 cilia. Asterisks indicate statistical significance value calculated by one-way ANOVA followed by Tukey’s post hoc test. ****, *p* ≤ 0.0001.

**Figure 6.**
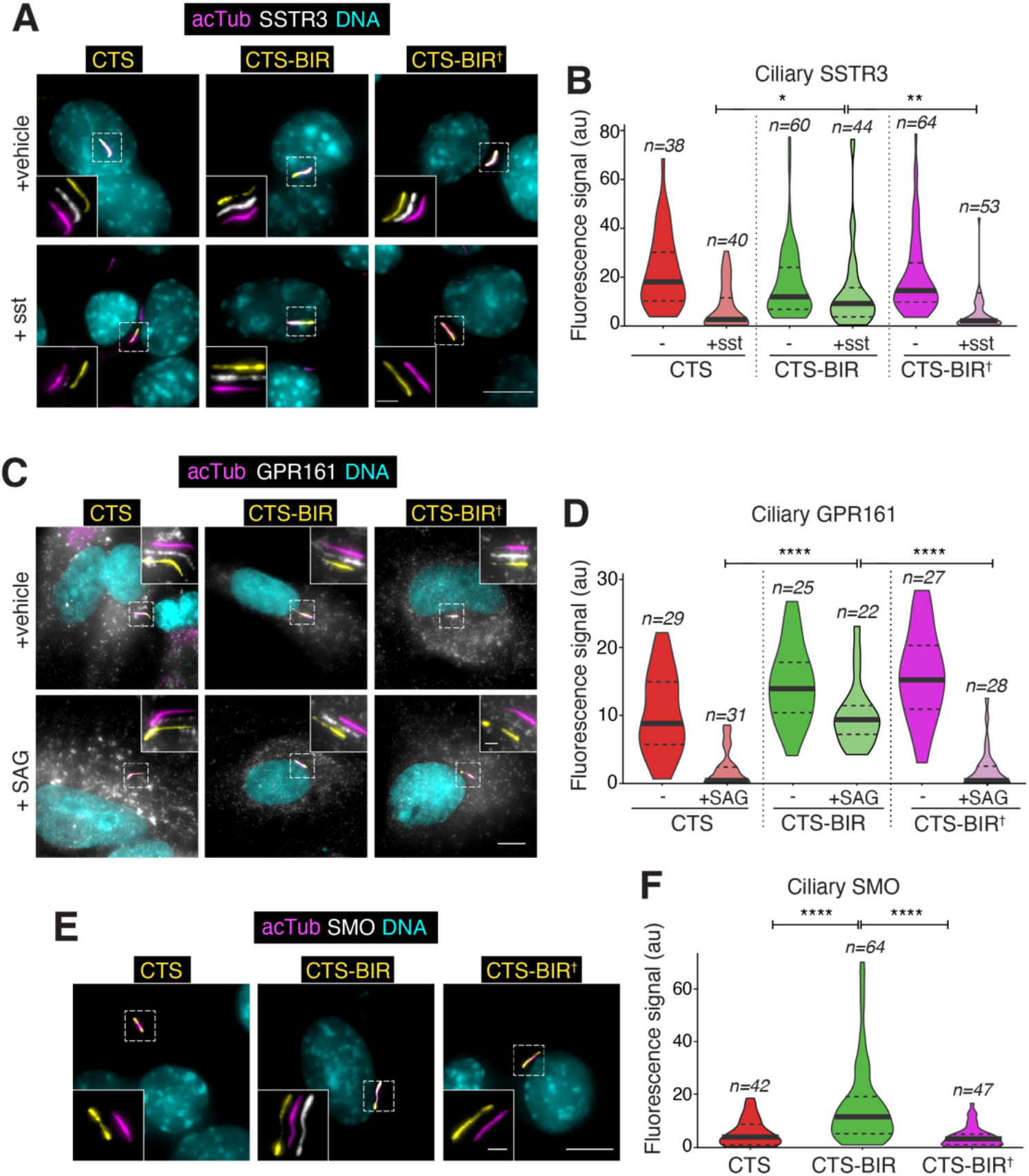
Targeted disruption of the BBSome-TOM1L2 interaction blocks the regulated exit of GPCRs from cilia. **A.** IMCD3-[pEF1 α^Δ^-^AP^SSTR3; pEF1α-BirA•ER] were transfected with plasmids expressing the BBSome interacting region of TOM1L2 (BIR) fused to the ciliary targeting signal of NPHP3 (CTS) and GFP or indicated variants. Ciliary ^AP^SSTR3 was pulse-labeled with Alexa647-labeled monovalent streptavidin (mSA647) for 5 to 10 min, and cells were then treated with or without sst for 2 h, before fixation and staining for acetylated tubulin (acTub; magenta) and DNA (cyan). The CTS fusions were visualized through the intrinsic fluorescence of GFP (yellow) and ^AP^SSTR3 was visualized via mSA647 (white). Scale bars, 5 μm (main panel), 1 μm (inset). **B.** The fluorescence intensities of ciliary ^AP^SSTR3 are represented as violin plots. Asterisks indicate statistical significance value calculated by one-way ANOVA followed by Tukey’s post hoc test. **, *p ≤* 0.01; *, *p ≤* 0.05. *n* = 38-64 cilia. **C.** RPE1-hTERT cells transfected with the indicated constructs were treated with SAG or vehicle (DMSO) for 2 h and then fixed and stained for acetylated tubulin (magenta), GPR161 (white), and DNA (cyan). The CTS fusions were visualized through the intrinsic fluorescence of GFP (yellow). Scale bars, 5 μm (main panel), 1 μm (inset). **D.** The fluorescence intensities of ciliary GPR161 are represented as violin plots. Asterisks indicate ANOVA significance value. ****, *p* ≤ 0.0001. **E.** IMCD3-[pCrys-SMO^FLAG^] cells transfected with the indicated constructs were fixed and stained for acetylated tubulin (magenta), FLAG (SMO, white), and DNA (cyan). The CTS fusions were visualized through the intrinsic fluorescence of GFP (yellow). **F.** The fluorescence intensities of ciliary SMO are represented as violin plots. Asterisks indicate ANOVA significance value. ****, *p* ≤ 0.0001. n = 42-64 cilia.

**Figure 7.**
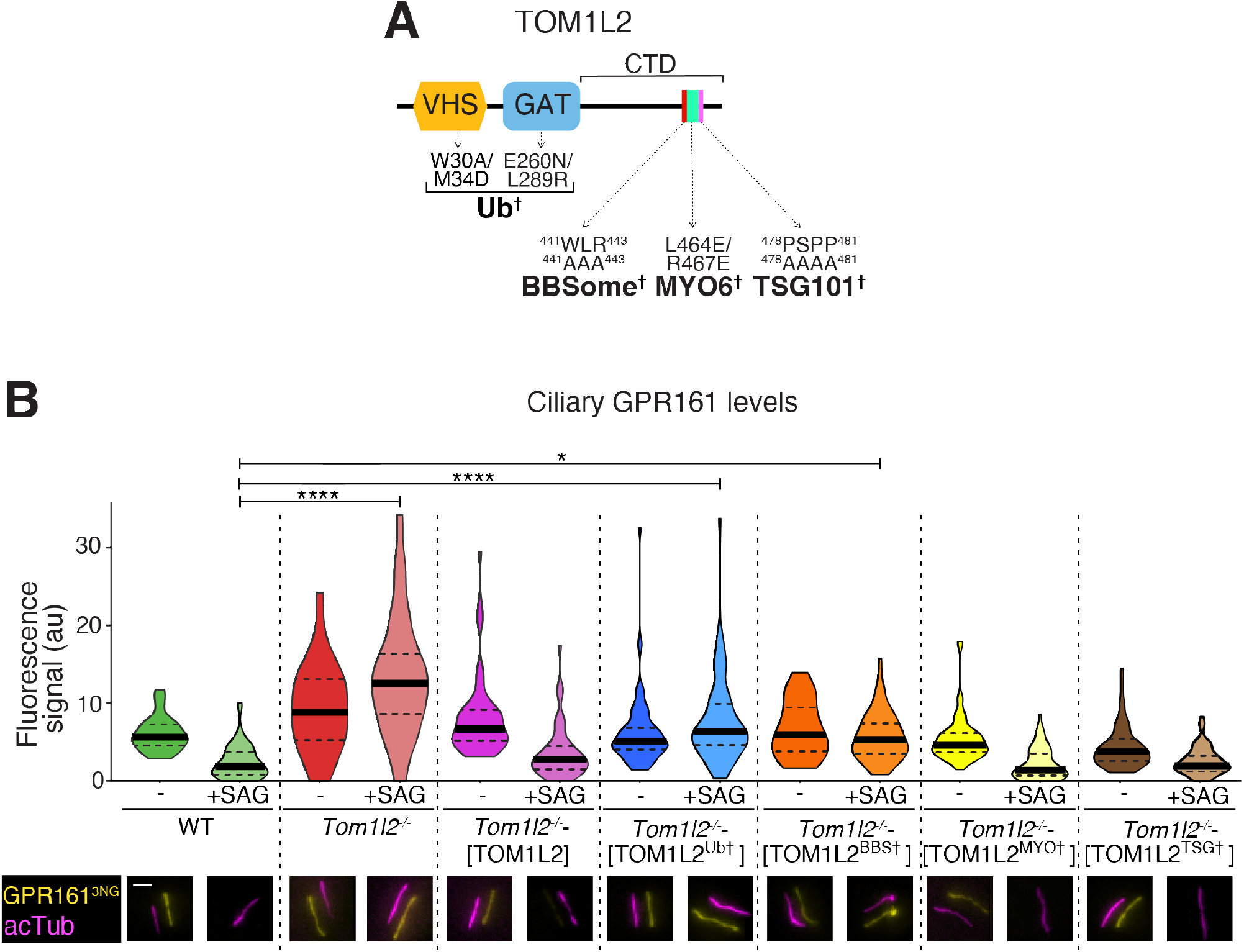
Rescue of *Tom1l2*^−/−^ cells demonstrates dual requirements for BBSome and Ub binding in supporting GPCR exit. **A.** Diagram of the domain organization of TOM1L2 with point mutations shown to disrupt interactions with the indicated partners. See **Fig. S8A-B** for variant validation **B.** TOM1L2 variants defective in interactions with known partners were stably expressed into *Tom1l2*^−/−^ IMCD3 cells together with GPR161^NG3^. In wildtype IMCD3 cells, only GPR161^NG3^ was stably expressed. Cells were treated with SAG or vehicle (DMSO) for 3 h and then fixed and stained for acetylated tubulin (magenta). GPR161^NG3^ was detected via the intrinsic fluorescence of NeonGreen. Scale bar, 1 μm. The fluorescence intensities of ciliary GPR161^NG3^ are represented as violin plots. Asterisks indicate statistical significance value calculated by one-way ANOVA followed by Tukey’s post hoc test. ****, *p* ≤ 0.0001. *, *p* ≤ 0.05.

### TOM1L2 acts downstream of the BBSome and upstream of Ub chain recognition

We next sought to determine how TOM1L2 sorts GPCRs out of cilia. TOM1L2 may bridge cargo-linked ubiquitin chains to retrograde BBSome trains or TOM1L2 may facilitate the movement of cargo-laden BBSome trains out of cilia. The latter mechanism is exemplified by BBSome regulators such as IFT27 or LZTFL1, whose depletion result in the drastic accumulation of BBSomes and associated cargoes inside cilia (Eguether et al., 2014; Liew et al., 2014). Meanwhile the ubiquitin bridging mechanism predicts that TOM1L2 is required for cargo exit but dispensable for BBSome exit. Indeed, removal of UbK63 chains from cilia via the K63-specific deubiquitinase AMSH fused to a ciliary targeting signal (CTS) blocks GPCR exit from cilia (Desai et al., 2020; Shinde et al., 2020) without affecting BBSome distribution inside cilia (**Fig. 3A and S6A**). A catalytically inactive AMSH does not affect GPCR exit from cilia and served as negative control (**Fig. 3A and S6A**). Similarly, depletion of TOM1L2 left BBSome distribution inside cilia unaffected (**Fig. 3C and S6C**) while interrupting GPCR exit. These data support a role of TOM1L2 as an adaptor between UbK63 chains and the BBSome.

In addition, we previously observed that activation of the Hh pathway via the Smoothened agonist SAG leads to a redistribution of BBSomes and its cargoes to the tip of cilia and surmised that this tip accumulation reflects a kinetically slow step of cargo loading onto departing BBSome trains (Ye et al., 2018). While BBSome tip accumulation was readily detected in cells transfected with catalytically inactive CTS-AMSH, the magnitude of the tip accumulation was greatly reduced in cells expressing CTS-AMSH (**Fig. 3B**). These data suggest that weakening of cargo attachment to BBSome upon removal of UbK63 chains leads to a decreased residence of BBSome at the tip of cilia because of an accelerated departure of BBSomes from the tip. Remarkably, depletion of TOM1L2 greatly reduced he BBSome tip accumulation observed upon Hh pathway activation, again phenocopying expression of CTS-AMSH (**Fig. 3D**). We conclude that TOM1L2 acts between the recognition of ubiquitin chains attached to cargoes and BBSome-mediated exit.

### TOM1L2 directly associates with the BBSome

The endosomal sorting function of ESCRT-0 proteins is accomplished by combining UbK63 chain recognition via VHS, GAT and UIM domains with binding to ESCRT-I (Schöneberg et al., 2017; Vietri et al., 2020). TOM1 family members combine conserved VHS and GAT domains at their N-terminus with more divergent C-terminal domains that contact the ESCRT-I subunit TSG101 as well as MYO6 and clathrin (Roach et al., 2021) (**Fig. 4A**). To test whether TOM1L2 directly interacts with the BBSome, we expressed GST fusions with TOM1L2 (and TOM1 or TOM1L1 as controls) and conducted capture assays of BBSome purified to near-homogeneity from bovine retina. Both TOM1L2 and TOM1 efficiently captured the BBSome while TOM1L1 did not (**Fig. 4B and S7A**). Deletion mapping revealed that BBSome recognition is encoded within the C-terminal domain (CTD) of TOM1L2 (**Fig. 4C and S7B**). The BBSome binding to TOM1 and TOM1L2 but not TOM1L1 is consistent with the 65 % sequence similarity between the CTDs of TOM1 and TOM1L2 and the significance divergence of the CTDs between TOM1L1 and either TOM1 or TOM1L2 (<20% similarity for the two pairwise comparisons). Further deletion mapping narrowed down the interacting region to a stretch of 73 amino acids (384-456, **Fig. 4D and S7C**) that also contains the binding determinants for MYO6 (Hu et al., 2019). Given that MYO6 accumulates in cilia when BBSome function is compromised and participates in ectocytosis (Nager et al., 2017), we sought to test whether MYO6 and the BBSome engage the same binding site on TOM1L2. We narrowed down the BBSome-binding region to a 30 amino acid segment in TOM1L2 CTD (384-413, **Fig. 4E and S7D**). As this minimal BBSome-binding fragment no longer binds to MYO6 (**Fig. S8A**), we conclude that TOM1L2 contains separable binding sites for the BBSome and MYO6.

To independently validate our deletion mapping, we modified the visual capture assay (Katoh et al., 2015) using fragments of TOM1L2 immobilized on beads to capture GFP-tagged human BBSomes expressed in HEK cells. Results of the visual capture assay were fully congruent with the capture of pure retinal BBSome (**Fig. 4F**), confirming that the minimal BBSome interacting region (BIR) of TOM1L2 comprises aa 384-413. To identify the amino acids responsible for BBSome binding inside TOM1L2, we conducted alanine scan mutagenesis on the BIR. A clear signal was detected within the first 10 amino acids of the BIR (**Fig. 4G**), pointing to the ^388^EVWLR^392^ motif as a BBSome-binding motif (BBM) in TOM1L2. Mutations in the BBM did not affect binding of TOM1L2 to MYO6 (**Fig. S8B**) and deletion of the entire BBM abolished binding of TOM1L2 to the BBSome (**Fig. S7E**) but not to MYO6 (**Fig. S8A**). These results further confirm that BBSome binding and MYO6 binding are separable on TOM1L2. Given that a 389-456 truncation retained some residual binding to the BBSome (**Fig. 4F**) and given that mutation of ^386^DIE^388^ to AAA only partially reduced binding, we conclude that ^389^VWLR^392^ defines a minimal BBSome-binding motif in TOM1L2. In TOM1, which also binds the BBSome, this motif is QWLS. This suggest that a tryptophan/leucine dipeptide may constitute the BBSome binding motif common to TOM1 and TOM1L2.

### TOM1L2 bridges the BBSome to ubiquitinated GPCR cargoes

The definition of a BBSome-binding motif in TOM1L2 enabled us to directly test the model that TOM1L2 functions as an adaptor between the BBSome and UbK63-linked proteins destined for ciliary exit (**Fig. 5A**). We reasoned that a cilia-localized BIR peptide will outcompete the BBSome-TOM1L2 interaction and result in the accumulation of TOM1L2 and ubiquitinated GPCRs inside cilia. The ciliary targeting signal (CTS) was NPHP3[1-200] and the ^390^WLR^392^ to AAA mutant of TOM1L2’s BIR (BIR^†^) provided a negative control for the experiment. While expression of the CTS either alone or fused to the BIR^†^ mutant left ciliary TOM1L2 levels barely detectable, expression of the CTS-BIR drastically and significantly increased the ciliary levels of TOM1L2 (**Fig. 5B-C**). Meanwhile, the BBSome distribution inside cilia was left largely unchanged by expression of CTS-BIR (**Fig. 5D-E and S6D**), thus indicating that disruption of the TOM1L2-BBSome interaction blocks TOM1L2 exit but leaves BBSome trafficking intact. Nonetheless, as we observed with expression of CTS-AMSH or depletion of TOM1L2 (**Fig. 3**), the tip enrichment of BBSome upon SAG addition was dampened in cells that expressed CTS-BIR (**Fig. 5E**). This decreased residence of BBSome at the tip of cilia indicates that disrupting the BBSome-TOM1L2 interaction prevents the efficient engagement of BBSome onto its cargoes at the tip of cilia.

We next tested for the importance of the BBSome-TOM1L2 interaction in ferrying ubiquitinated proteins out of cilia. In SAG-treated cells where GPR161 undergoes stimulated exit, CTS-BIR expression led to a considerable and significant increase in the ciliary levels of ubiquitin compared to expression of CTS alone or CTS-BIR† (**Fig. 5F-G**). These data show that the BBSome needs to recruit TOM1L2 in order to remove ubiquitinated proteins from cilia.

For the final test of our model, we monitored the ciliary exit of the BBSome cargoes SMO, GPR161 and SSTR3. The ciliary dynamics of SSTR3 and SMO were assayed in IMCD3 cells stably expressing tagged GPCRs and the exit of endogenous GPR161 was assessed in RPE1-hTERT cells. While SSTR3 underwent sst-dependent exit from cilia in cells transfected with the controls CTS and CTS-BIR†, expression of CTS-BIR blocked the ciliary exit of SSTR3 (**Fig. 6A-B**). Similarly, GPR161 exit upon Hedgehog pathway activation proceeded normally in RPE cells transfected with CTS or and CTS-BIR†, but expression of CTS-BIR considerably blunted the ciliary exit of GPR161 (**Fig. 6C-D**). Finally, expression of CTS-BIR was sufficient to promote the accumulation of SMO in cilia in the absence of pathway activation. In contrast, expression of the controls CTS and CTS-BIR^†^ left the ciliary levels of SMO nearly undetectable, strongly suggesting that the CTS-BIR fusion blocks the constitutive ciliary exit of SMO (**Fig. 6E-F**). We conclude that disruption of the BBSome-TOM1L2 interaction blocks the regulated exit of GPCRs from cilia. Together, these results demonstrate that ubiquitinated GPCRs are recognized inside cilia by TOM1L2 before latching onto the BBSome/IFT exit machinery and undergoing removal from cilia.

### TOM1L2 binding to Ub and BBSome is required for ciliary retrieval

Given that TOM1L2 interacts with TSG101, MYO6, BBSome and Ub, we sought out to test which of these interactions was essential to the function of TOM1L2 in retrieval of ubiquitinated GPCRs from cilia. Knockout of TOM1L2 in IMCD3 cells completely blocked the signal-dependent exit of GPR161 from cilia (**Fig. 7B**), in agreement with the siRNA data obtained in IMCD3 cells (**Fig. 2D and S4**). Crystallographic studies have mapped the binding sites for MYO6 and ubiquitin on TOM1L2 (Akutsu et al., 2005; Hu et al., 2019) (**Fig. 7A**) and we validated point mutants defective in binding to MYO6 (**Fig. S9A**) or UbK63 chains (**Fig. S9B**).

TSG101 has been shown to interact with TOM1L1 via the known TSG101-binding motif P[S/T]xP (PSAP and PTAP in TOM1L1) (Yanagida-Ishizaki et al., 2008) and TOM1L2 contains a P[S/T]xP motif (PSPP in TOM1L2), which we mutated to AAAA. Measurement of GPR161 signal-dependent exit in stable cell lines reexpressing the TOM1L2 variants demonstrated a requirement for the interactions of TOM1L2 with Ub and BBSome (**Fig. 7B**). Meanwhile, the interactions of TOM1L2 with MYO6 and TSG101 were dispensable for the retrieval of GPR161 from cilia. These data confirm that TOM1L2 acts as an adaptor between the BBSome and ubiquitinated GPCRs inside cilia and that neither MYO6 nor TSG101 assist TOM1L2 in the retrieval process.

### Evolutionary conservation of TOM1L2 in ciliary clearance of ubiquitinated proteins

To determine whether TOM1L2’s function in ciliary exit is conserved outside of mammals, we turned our attention to the single cell free-living organism *Chlamydomonas reihnardtii*. Humans are separated from *Chlamydomonas* by nearly 500 million years of divergent evolution, and *Chlamydomonas* is a well-validated system for ciliary research in general and BBSome-mediated exit in particular (Lechtreck et al., 2009, 2013; Liu and Lechtreck, 2018; Xue et al., 2020). Phylogenetic searches identified the gene *Cre06.g292000* as the single representative of the TOM1 family in the *Chlamydomonas* genome (Herman et al., 2011). *Chlamydomonas* TOM1 contains GAT and VHS domains and reciprocal BLAST of *Chlamydomonas* TOM1 against the human genome returns TOM1L2 as the top hit. Several *tom1* mutants were recovered in the CLiP library of insertional *Chlamydomonas* mutants (Li et al., 2019) and two of them are predicted to be complete loss of function alleles.

Staining of the two *tom1* mutants for ubiquitin revealed a significant enrichment of ubiquitin in cilia compared to the wildtype strain (**Fig. 8A-B**), as previously reported in *bbs4* mutants (Shinde et al., 2020). The role of TOM1L2 in removing ubiqutinated proteins from cilia thus appears to be conserved in *Chlamydomonas*. We next turned our attention to phospholipase D (PLD), a well-validated BBSome cargo that is normally undetectable in cilia but markedly accumulates in cilia of *bbs* mutants (Lechtreck et al., 2013). Surprisingly, while PLD accumulation was readily detected in cilia of *bbs4* mutants, no such accumulation was detected in *tom1* mutant cilia (**Fig. 8C-D**). One possible interpretation is that PLD represents a BBSome cargo that does not require ubiquitination or crTOM1 for its exit. In this context, it is worth noting that there is currently no evidence that PLD is ubiquitinated in cilia of *bbs* mutants as no high molecular weight smear indicative of ubiquitination is detected when immunoblotting the ciliary pool of PLD in *bbs* mutants (Lechtreck et al., 2013). Nonetheless, care should be exercised when interpreting such results as highly active deubiquitinases will rapidly digest Ub chain upon cell lysis unless specific inhibitors are included in the lysis buffer (Hjerpe et al., 2009). The status of PLD ubiquitination in cilia and the role of ubiquitination in PLD exit will thus need to be resolved by future experimentation.

**Figure 8.**
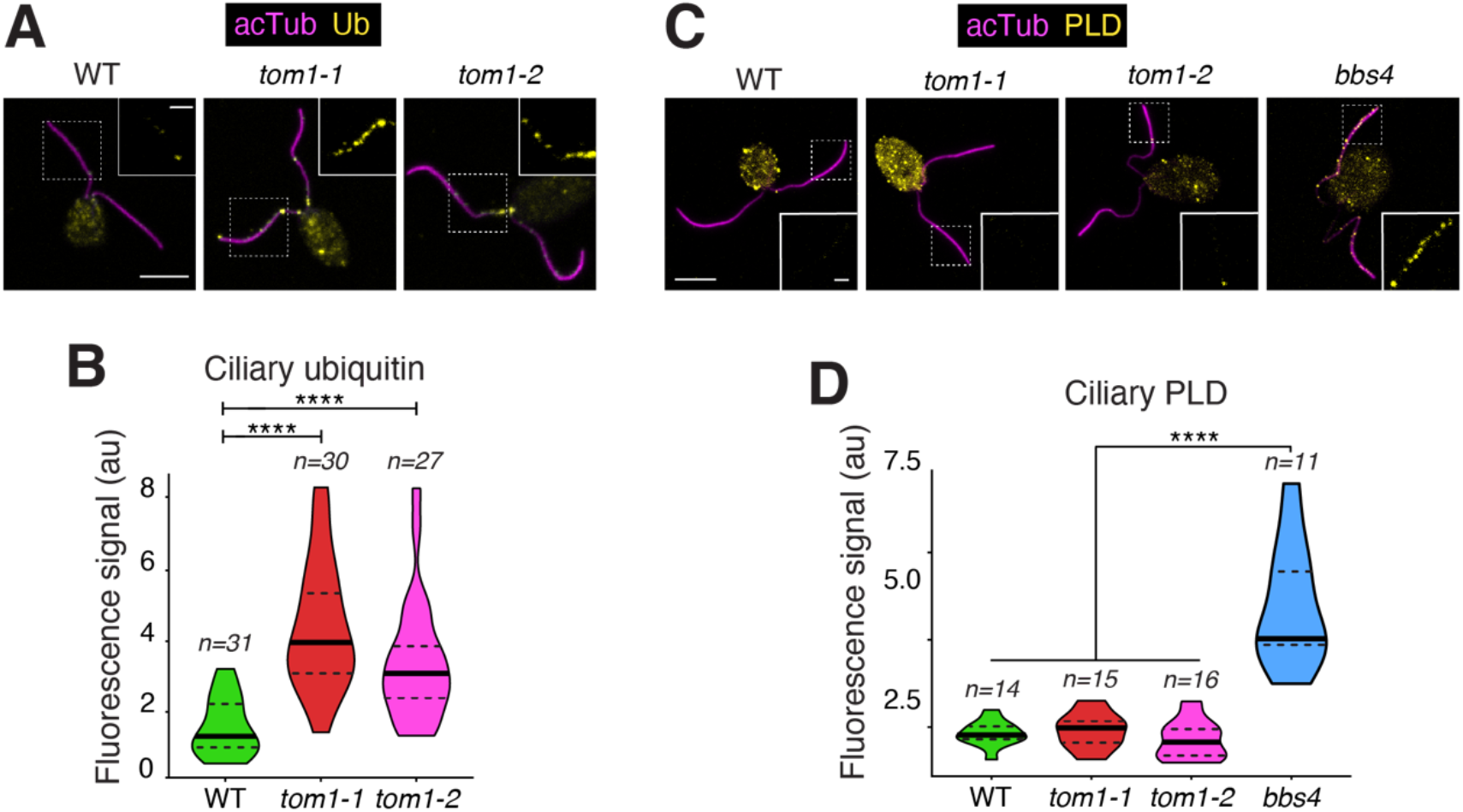
*Chlamydomonas rheinardtii* TOM1 is required for the removal of ubiquitinated proteins from cilia. **A.** WT or *tom1* mutant *C. rheinardtii* cells were fixed and stained for acetylated tubulin (acTub, magenta) and Ub (yellow). Scale bars, 5 μm (main panel) and 1 μm (inset). **B.** Violin plots of the ciliary Ub levels in WT and *tom1 C. rheinardtii* cells. Asterisks indicate statistical significance value calculated by one-way ANOVA followed by Dunnet’s post hoc test. ****, *p ≤* 0.0001. The ciliary levels of Ub are increased more than three to four-fold in *tom1* cells as compared with WT. **C.** *C. rheinardtii* cells of indicated genotypes were fixed and stained for acetylated tubulin (acTub, magenta) and PLD (yellow). Scale bars, 5 μm (main panel) and 1 μm (inset). **D.** Violin plots of the ciliary PLD levels in WT or *tom1* or *bbs4 C. rheinardtii* cells. Asterisks indicate statistical significance value calculated by one-way ANOVA followed by Dunnet’s post hoc test. ****, *p* ≤ 0.0001. The ciliary levels of PLD are increased more than 2.5 fold in *bbs4* cells compared to other genotypes.

## DISCUSSION

### TOM1L2 is a conserved adaptor for the retrieval of UbK63-tagged proteins from cilia

Unbiased proteomics profiling of *Bbs* cilia and a focused screen of ESCRT proteins and related UbK63 readers identified TOM1L2 as a ciliary protein that is required for the regulated exit of GPCRs from cilia. Targeted disruption of the BBSome-TOM1L2 interaction and functional rescue with point mutants of TOM1L2 reveals that TOM1L2 bridges ubiquitinated GPCRs to the retrieval machinery. Finally, the role of TOM1L2 in sorting ubiquitinated proteins out of cilia is conserved in *Chlamydomonas*. We conclude that TOM1L2 is a conserved adaptor that enables the ciliary retrieval machinery to recognize and ferry cargoes marked by UbK63 chains. The role of TOM1L2 in ciliary biology is independently suggested by findings of canonical ciliopathy symptoms in a mouse model of *Tom1l2* deficiency (Girirajan et al., 2008). Besides a high incidence of infections and tumors, mice homozygous for a hypomorphic *Tom1l2* allele show high incidence of tooth malocclusion, kyphosis, hydrocephaly, and renal cysts.

We note that no binding of IFT-A, IFT-B or BBSome to UbK63 chains was detected in our experimental systems. In contrast, Wang and colleagues found that the IFT-A subunit IFT139 captures ubiquitinated proteins (including α-tubulin) from ciliary extracts of *Chlamydomonas* kinesin II mutants (Wang et al., 2019). It is conceivable that species-specific differences may exist in how ubiquitinated proteins are recognized by the ciliary retrieval machinery, or that IFT139 recognizes some ciliary proteins that happen to get ubiquitinated in kinesin II mutants. Direct interaction assays between UbK63 chains and the purified IFT-A complex will be needed to distinguish these hypotheses.

### TOM1 family proteins are conserved ubiquitin adaptors for the autophagy, endosomal and ciliary machineries

In plants, transporters and carrier proteins undergo regulated degradation in response to environmental changes via sorting from the plasma membrane to the vacuole, the plant equivalent of the lysosome. Most aspects of endolysosomal sorting are conserved between mammalian and plant systems with ubiquitination, ESCRT-I, -II and -III governing the sorting of membrane proteins to the vacuole in plants (Gao et al., 2017). Yet, HRS and STAM homologues are only found in opisthokonts (i.e. fungi and animals), and absent from amoeba, plants and flagellated single cell organisms (Herman et al., 2011; Leung et al., 2008). The absence of the canonical ESCRT-0 HRS/STAM in plants remained a puzzling exception to the conserved function of ESCRT complexes until TOM1 homologues were identified in every eukaryotic branch and found to play roles in endolysosomal sorting in plants and mammalian systems (Mosesso et al., 2019). In the model plant *Arabidopsis thaliana*, the combined ablation of five out of nine TOM1-like proteins (TOLs) interrupts the sorting of the transmembrane auxin carrier protein PIN2 (Korbei et al., 2013) and of the boron transporter BOR1 (Yoshinari et al., 2018) at the level of the plasma membrane and endosomes. Both PIN2 and BOR1 are ubiquitinated under conditions that promote their degradation (Kasai et al., 2011; Leitner et al., 2012) and the penta-TOL mutant accumulates considerable levels of ubiquitinated proteins (Moulinier-Anzola et al., 2020). Finally, nearly all studied TOLs have been detected on early endosomes (Korbei et al., 2013; Moulinier-Anzola et al., 2020; Yoshinari et al., 2018). Together, these data indicate that TOLs sort ubiquitinated membrane proteins to the degradative endolysosomal pathway.

A role of TOM1 in endosomal sorting in mammalian systems is evidenced by the signal-dependent accumulation of the Interleukin 1 receptor (IL1R) in late endosomes when TOM1 is knocked down in MEFs (Brissoni et al., 2006), and the requirement for TOM1 in signal-dependent degradation –but not internalization– of the delta opioid receptor (DOR) (Lobingier et al., 2017). Together, these data argue that TOM1 family proteins represent the ancestral component of the ESCRT-0 machinery and that the canonical ESCRT-0 HRS/STAM is a more recent evolutionary elaboration in fungi and animals.

Besides endolysosomal sorting and ciliary exit, UbK63 chains also function in autophagy by marking aggregates, organelles and pathogens for engulfment into autophagosomes (Grumati and Dikic, 2018). The defect in autophagosome maturation in cells depleted of TOM1 and TOM1L2 and the localization of TOM1 to autophagosomes indicate that TOM1 may directly participate in autophagy (Tumbarello et al., 2012). The conserved and diverse roles of TOM1 family proteins in sorting of UbK63-marked cargoes befits the ancestral origin of TOM1 and supports a universal role for TOM1 family members in the first step of UbK63-marked cargo sorting.

### Post-exit fate of ciliary GPCRs

In plants, TOM1 family proteins appear to initiate recognition of their cargoes at the plasma membrane. TOL6 localizes to the plasma membrane of *Arabidopsis* root cells at steady state (Moulinier-Anzola et al., 2020), the pentaTOL *Arabidopsis* mutant accumulates a PIN2-Ub fusion at the plasma membrane (Korbei et al., 2013) and UbK63 is detected at the plasma membrane, as well as endosomes and vacuoles (Johnson and Vert, 2016). TOM1 family proteins thus appear to recognize their ubiquitinated cargoes early and escort them from the plasma membrane to late endosomes. In this context, TOM1L2 may escort ubiquitinated proteins from cilia to the lysosome, first by bridging them to the BBSome, then to the endocytic machinery and finally by transferring them to the ESCRT machinery for ultimate lysosomal degradation. Testing of this fascinating hypothesis awaits the development of TOM1L2 mutants that fail at supporting endocytosis and lysosomal sorting as well as techniques that can track GPCRs from cilia to the lysosome.

### A number of UbK63 readers are detected inside cilia of *Bbs* mutants

The accumulation of six UbK63 readers in cilia of retrieval mutants is surprising when one considers that only TOM1L2 was found to function in retrieval. What may be the roles of MYO6, TOM1, TOLLIP, HRS and STAM inside cilia? First, the biochemical interaction of TOM1 with the BBSome and the increase in TOM1 levels inside *Arl6* and *Ift27* cilia (Mick et al., 2015) and **Fig. 2A**) suggest that TOM1 may function redundantly with TOM1L2 in the retrieval of ubiquitinated GPCRs. However, any possible role for TOM1 in retrieval needs to be minor compared to TOM1L2 as no GPCR exit defect was detected in cells depleted of TOM1 and the double depletion of TOM1 and TOM1L2 did not further affect GPCR exit beyond depletion of TOM1L2 alone. Second, some UbK63 readers may participate in other ciliary trafficking modalities besides GPCR retrieval. In prior work, we uncovered a role for MYO6 in ectocytosis of GPCRs from cilia (Nager et al., 2017) and it is conceivable that ubiquitin and some Ub readers participate in ciliary ectocytosis. The very close proximity of the binding sites for BBSome and MYO6 on TOM1L2 raises the possibility that steric hindrance may prevent MYO6 and the BBSome to coincidently bind to TOM1L2. It will be important to test whether a possible competition between the BBSome and MYO6 for TOM1L2 binding may influence the decision to exit via retrieval vs. ectocytosis. Third, although neither HRS nor STAM scored as hits in our GPCR retrieval screen, they participate in the exit of ubiquitinated proteins out of nematode cilia (Hu et al., 2007) and may function in the retrieval of other cargoes besides GPCRs. Fourth, it is conceivable that the accumulation of UbK63 chains inside *bbs* cilia traps UbK63 readers inside cilia. Because only select UbK63 readers accumulate inside *bbs* cilia, we consider this broad hypothesis unlikely. It is nonetheless possible that TOLLIP and MYO6 accumulate in cilia when TOM1L2 is trapped inside cilia of *bbs* mutants because these two proteins directly associate with TOM1L2.

### Role of TOM1L2 in ciliary exit of constitutive vs. regulated cargoes

The increased Ub signal in cilia of *bbs* and *tom1* mutants compared to WT *Chlamydomonas* suggests that Ub marks some proteins for removal from *Chlamydomonas* cilia as it does for ciliary signaling receptors in mammals and nematodes. The nature of the ubiquitinated proteins that accumulate in *Chlamydomonas* cilia remains to be determined, in particular whether these represent signaling receptors that undergo regulated exit under vegetative growth conditions or whether they correspond to proteins that accidentally enter cilia and need to be constitutively removed from cilia. As imaging studies combined with genetics have extensively validated PLD as a BBSome cargo in *Chlamydomonas* and there is currently no indication that the ciliary localization of PLD responds to signaling inputs, PLD represents the paradigm of a constitutive BBSome cargo. In this context, the absence of PLD accumulation inside *Chlamydomonas tom1* mutant cilia suggests either that constitutive BBSome cargoes undergo Ub-independent exit from cilia or that additional adaptors besides TOM1 link ubiquitinated proteins to the BBSome. As noted above, some of the UbK63 readers that accumulate inside *Bbs* mutant cilia could function in the constitutive retrieval of non-ciliary proteins. Against this hypothesis, all other UbK63 readers that accumulate inside cilia of *Bbs* mutant IMCD3 cells besides TOM1/TOM1L2 are not conserved in *Chlamydomonas*. The hypothesis that constitutive exit is Ub-independent implies that PLD is recognized by the BBSome as a non-ciliary protein without the help of ubiquitin chains. In the future, it will be important to determine whether ubiquitin-independent cargoes of the BBSome do exist in *Chlamydomonas* and in mammalian systems. Constitutive removal of non-ciliary proteins has also been proposed to function in the clearance of pollutants that enter photoreceptor cilia (also known as outer segments) (Datta et al., 2015), and future work will need to test the role of ubiquitin in retrieval from the photoreceptor outer segment and the identity of the ubiquitin reader that may participate in this process. Regardless of the specific mode of PLD recognition by the BBSome, the constitutive removal of non-ciliary proteins from cilia poses the fascinating question of how proteins may be recognized as foreign to the cilium by the BBSome or by the ubiquitination machinery.

## ACKNOWLEDGEMENTS

We thank Folma Buss, Pamela Tran, Kinga Bujakowska, Hiroshi Hamada, Val Sheffield, Kathryn Anderson and Harald Stenmark for the gifts of antibodies; James Nathan, Rohan Baker, Folma Buss, Kirk Mykytyn, David Komander, Yuichiro Miyaoka and Greg Pazour for the gifts of plasmids; Karl Lechtreck for the gift of *Chlamydomonas* strains and the PLD antibody; the *Chlamydomonas* Mutant Library Group at Princeton University, the Carnegie Institution for Science, and the *Chlamydomonas* Resource Center at the University of Minnesota for providing the *Chlamydomonas tom1* insertional mutants; Yien-Ming Kuo for help with microscopy; Mingli Zhu for the gift of the stable IMCD3 cell line expressing SMO^Flag^; Stanford University Mass Spectrometry for data acquisition and all members of the Nachury lab for stimulating discussions. This work was funded by NIH (GM089933 and EY031462 to MVN; GM96745 to SPG) and ADA (1-20-VSN-03 to MVN). SRS acknowledges funding from the UCSF Program for Breakthrough Biomedical Research (7000/7002124) and International Retina Research Foundation and DUM from the Deutsche Forschungsgemeinschaft (TRR152/TP-25). This work was made possible, in part, by EY002162 - Core Grant for Vision Research and by the Research to Prevent Blindness Unrestricted Grant (MVN).

Proteomics data to be deposited in ProteomeXchange.

## MATERIALS AND METHODS

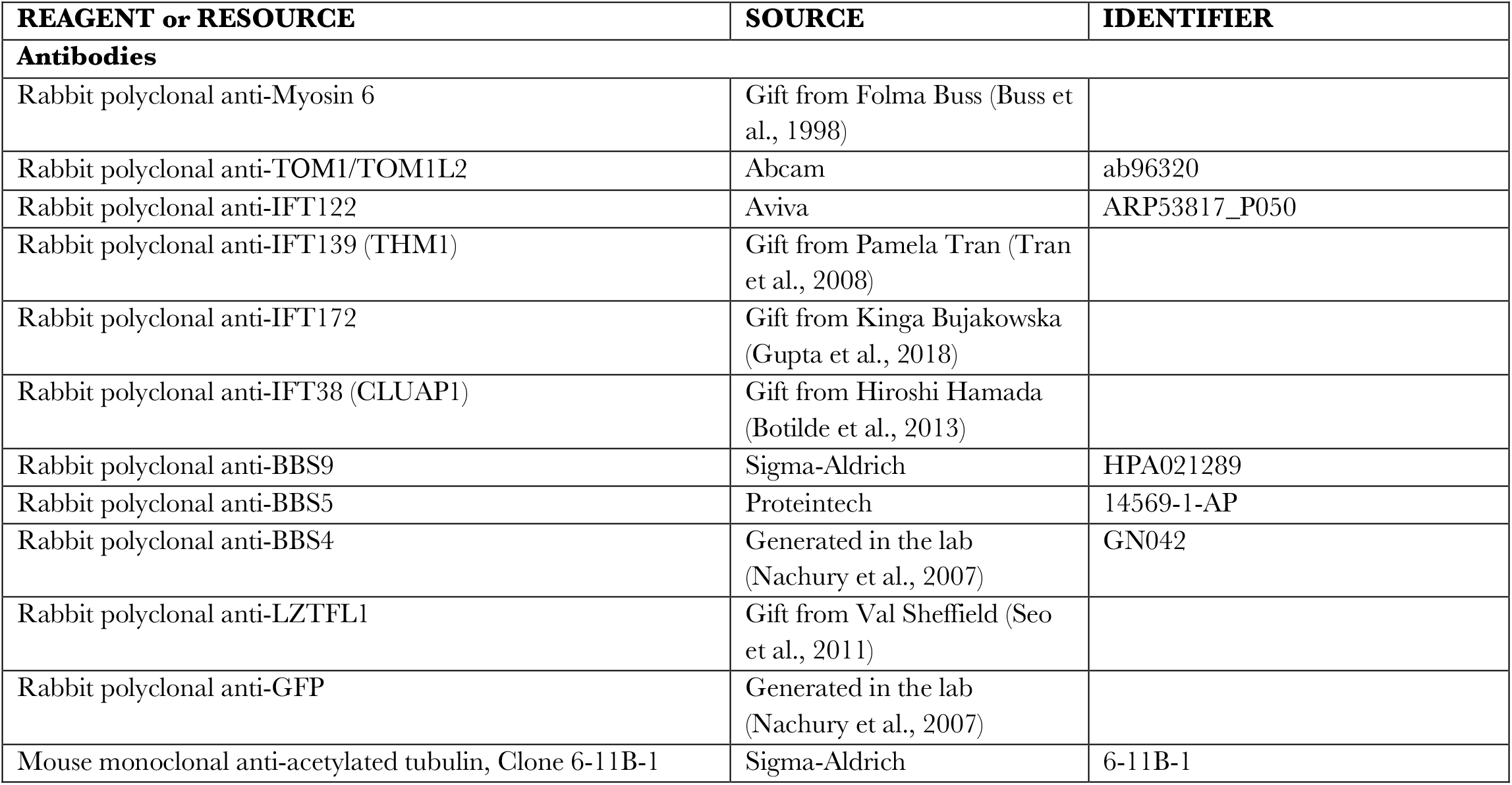
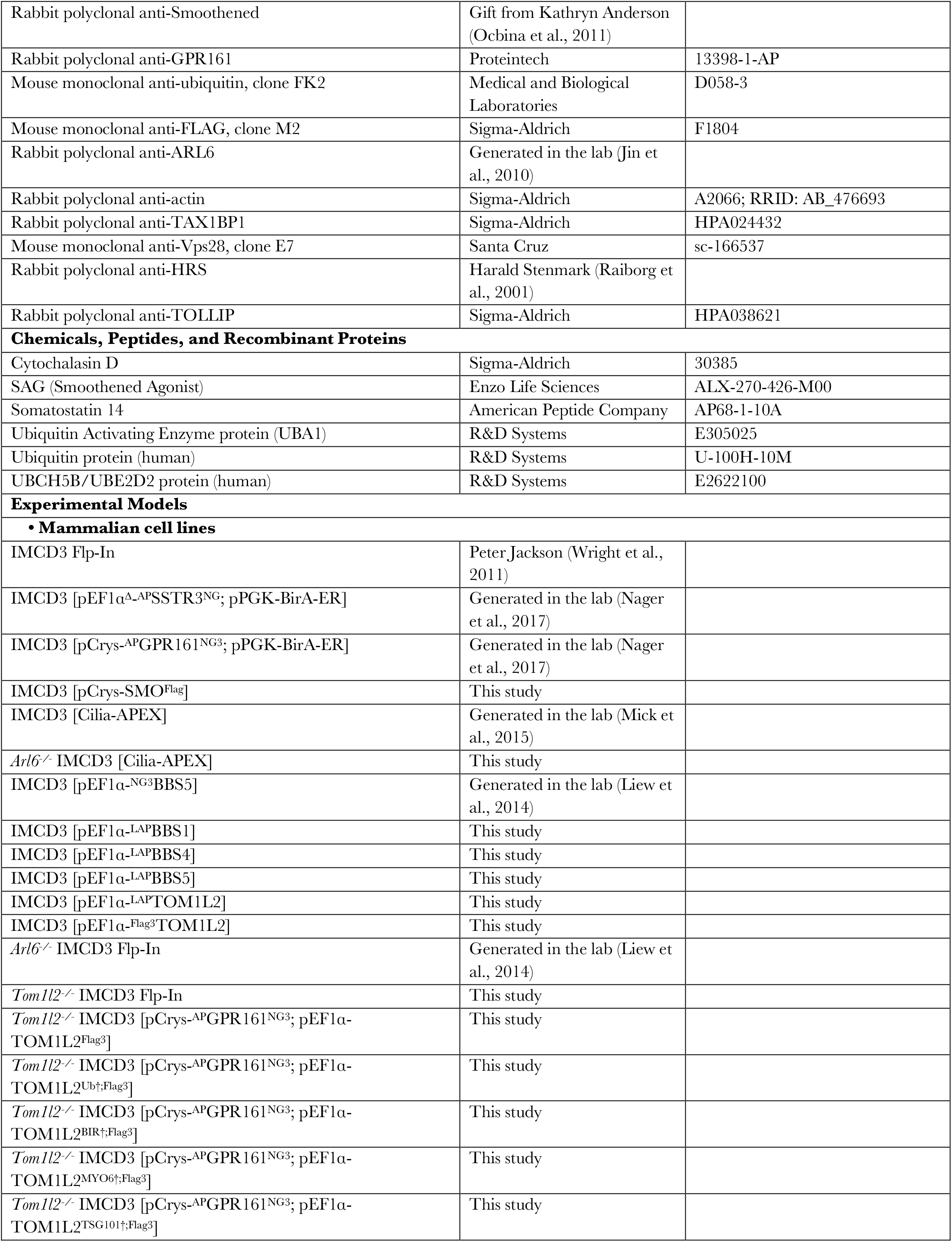
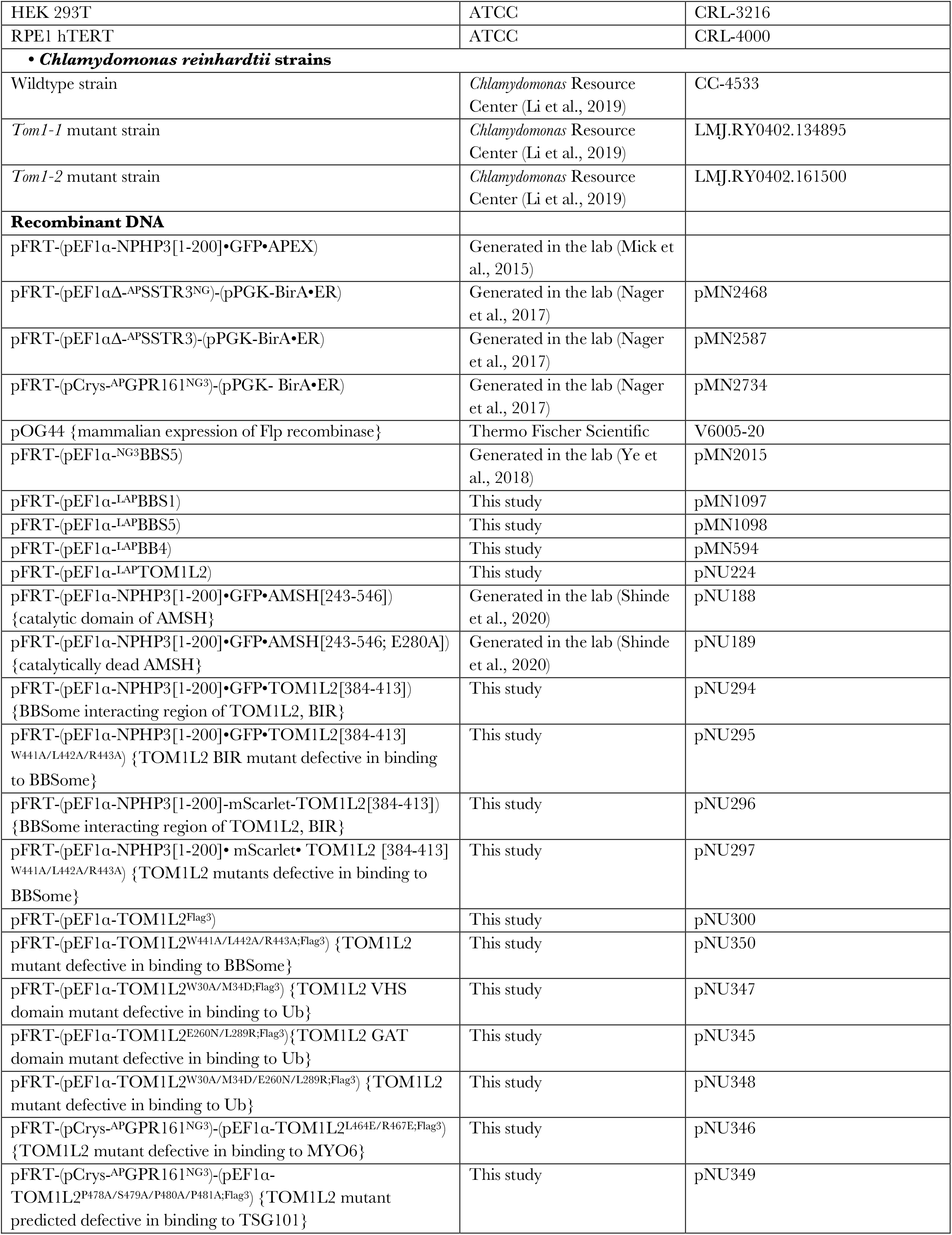
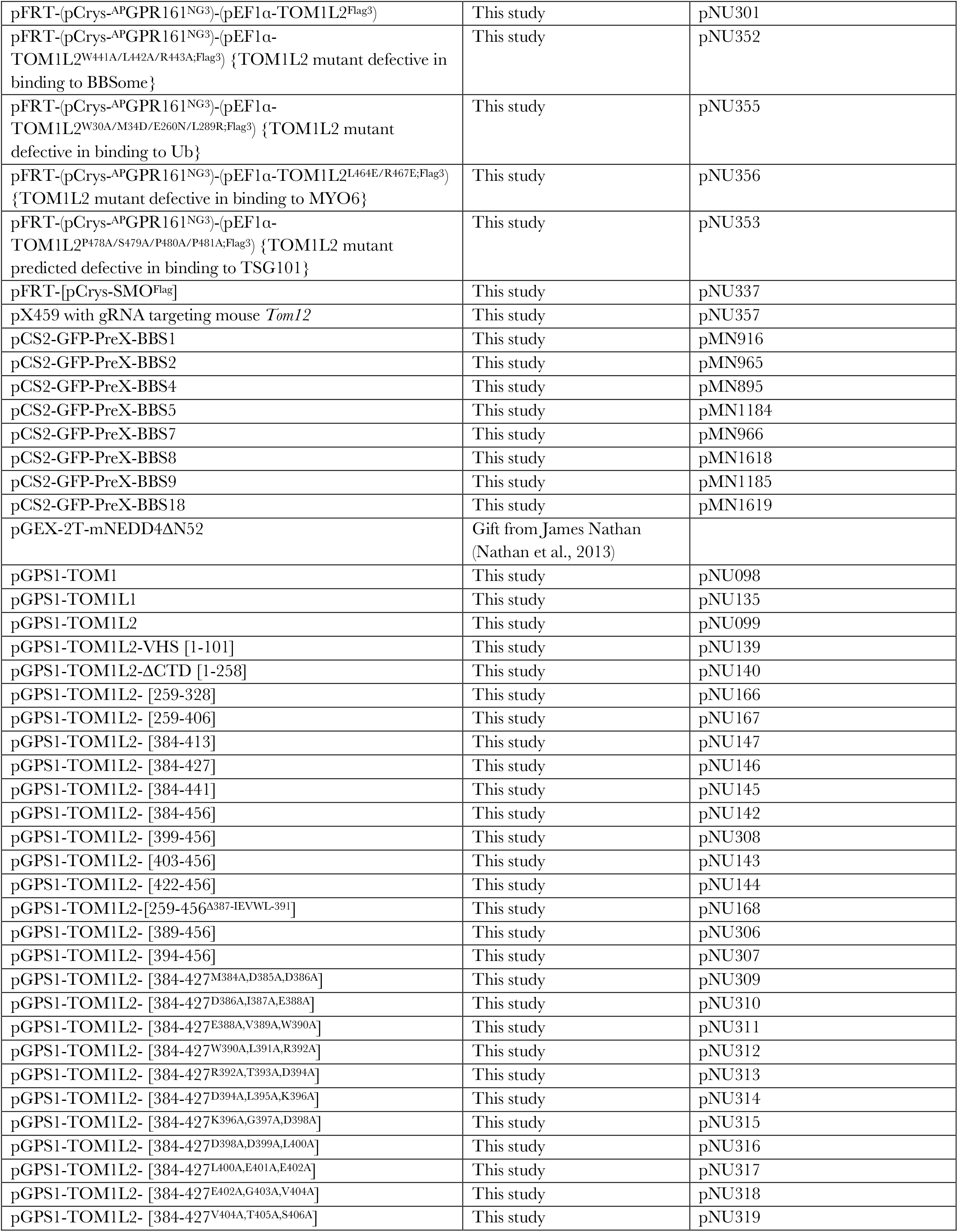
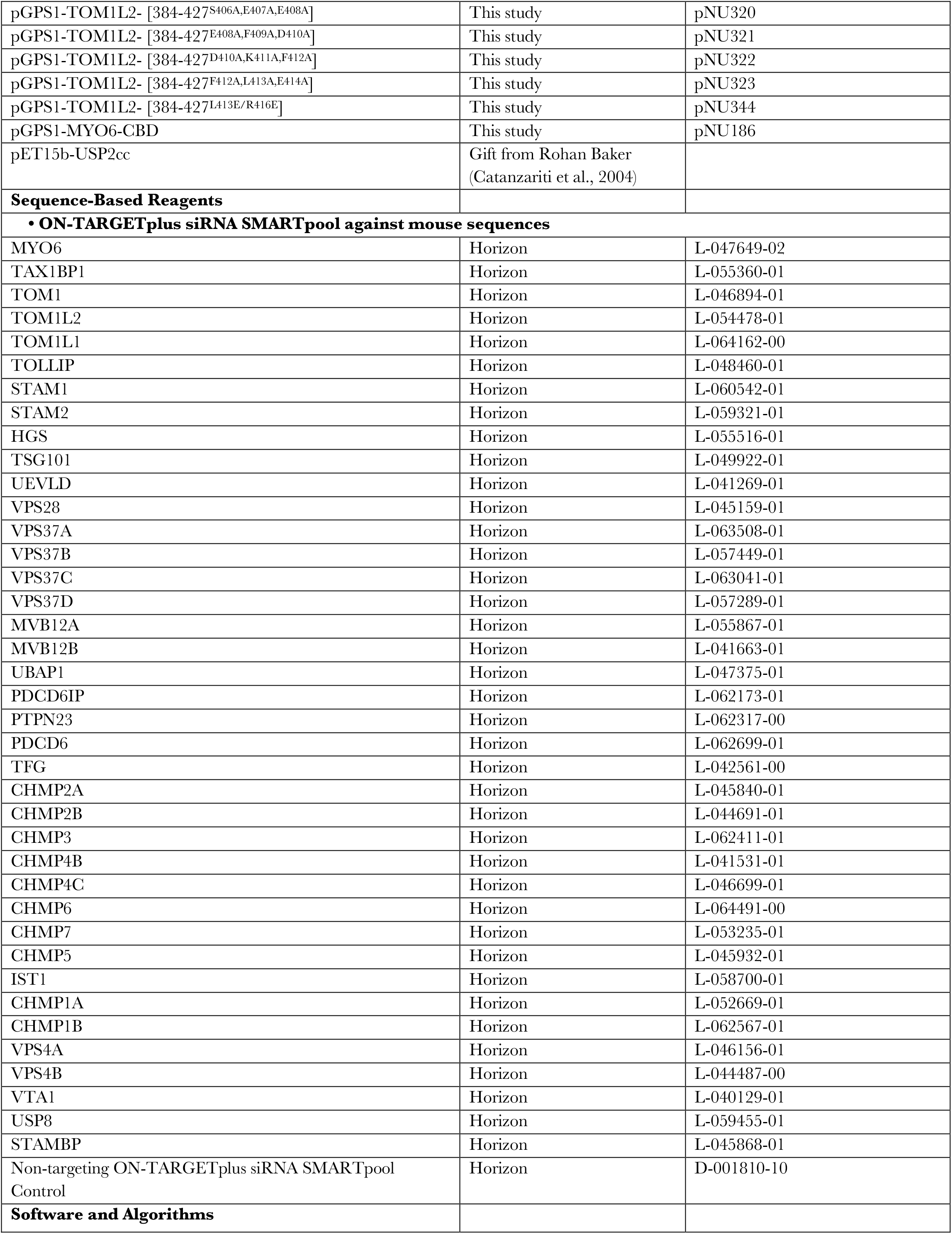
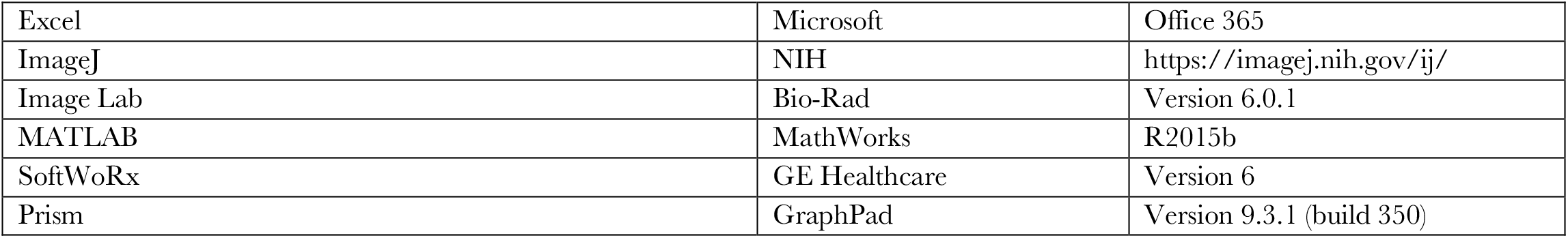

### RESOURCE AVAILABILITY

#### Lead contact

Further information and requests for resources and reagents should be directed to and will be fulfilled by the lead contact Maxence Nachury.

#### Materials availability

This study has generated plasmids and cell lines, which are listed in the Key resources table. These reagents will be made available upon request.

#### Data and code availability

All data reported in this paper will be shared by the lead contact upon request. This paper does not report original code. Any additional information required to reanalyze the data reported in this study is available from the lead contact upon request.

### EXPERIMENTAL MODEL AND SUBJECT DETAILS

All the IMCD3 cells used in the study were derived from a parental IMCD3-FlpIn cell line described previously (Jin et al., 2010). IMCD3 and RPE1-hTERT cells were cultured in DMEM/F12 supplemented with 10% FBS at 37°C with 5% CO_2_. Ciliation was induced by serum starving cells in media containing 0.2% FBS for 16 to 24 h.

*Chlamydomonas* cells were grown synchronously in a 14hour:10hour light:dark cycle in Tris-acetate-phosphate media (Gorman and Levine, 1965) for 72 h.

The Flp-In system (Thermo Fisher Scientific) was used to generate stable isogenic IMCD3 cell lines with single cassette integration. SSTR3, SMO and GPR161 fusion proteins were expressed at near-endogenous levels via attenuated promoters as described in (Ye et al., 2018) to reconstitute the appropriate ciliary trafficking dynamics.

### METHOD DETAILS

#### Plasmids

For bacterial expression, human TOM1 (Horizon MGC cDNA MHS6278-202760163), human TOM1L1 (Horizon MGC cDNA MHS6278-202840799) and human TOM1L2 (pGFP-TOM1L2, short isoform, gift from Folma Buss (Tumbarello et al., 2012)) were amplified and cloned into pGPS1 (GST-PreX-Stag-) by conventional cloning. pGSP1-TOM1L2 deletion domain mutants were generated by conventional cloning or by introducing a stop codon via site-directed mutagenesis. Site-directed mutagenesis or linker cloning was performed to generate the pGPS1-TOM1L2 AAA mutants. Human MYO6-CBD (aa 1060-1253) was amplified from pEGFP-MYO6, a gift from Folma Buss (Cambridge Institute for Medical Research, University of Cambridge, UK) (Buss et al., 1998), and cloned into pGPS1 by conventional cloning. pET15b-USP2cc was a kind gift from Rohan Baker (The Australia National University, Canberra, Australia) (Catanzariti et al., 2004). Mammalian expression constructs for GFP-tagged human BBSome subunits were generated by Gateway recombination of pCS2-GFP-Prex-DEST with pENTR-BBS1, BBS2, BBS4, BBS5, BBS7, BBS8, BBS9, or BBIP10. pGEX-2T-mNEDD4△N52 was a gift from James Nathan (University of Cambridge, Cambridge, UK). Mammalian expression vectors for LAP (localization and tandem affinity purification)-tagged BBS1, BBS4, BBS5, and TOM1L2 were generated by Gateway recombination of pEF5B•FRT•LAP-DEST with pENTR-BBS1, BBS4, BBS5, or TOM1L2. Coding sequences for human GPR161 (Horizon MGC cDNA MHS6278-202802001), mouse SSTR3 (gift from Kirk Mykytyn), mouse SMO (gift from Gregory Pazour, UMass, Worcester, USA; plasmid no. 164532; Addgene (Desai et al., 2020)), human BBS5 (gift from V. Sheffield, University of Iowa, Iowa City, IA) and human TOM1L2 (long isoform NP_001076437 gene synthesized by Genscript) were PCR amplified and cloned in FRT vectors with mNeonGreen (NG), birA Acceptor Peptide (AP) or the FLAG epitope. SSTR3 expression was driven by pEF1α^Δ^, GPR161 and SMO expression by pCrys, BBS5 and TOM1L2 expression by pEF1α. Cilia-targeted TOM1L2^BIR^ was expressed by fusing the BBSome interacting region of TOM1L2 (384-413) with NPHP3[1–200] and GFP or mScarlet to generate NPHP3[1-200]-GFP-TOM1L2[384-413] or NPHP3[1-200]-mScarlet-TOM1L2[384-413]. A BBSome interaction deficient version of TOM1L2 was generated by mutating WLR to AAA. Cilia-targeted AMSH was previously described (Shinde et al., 2020).

To knock out *TOM1L2* in IMCD3 cells, a gRNA targeting exon three of mouse *TOM1L2* (GCTCTAAAGAAGCGGCTTAG) was cloned into pX459V2.0-eSpCas9(1.1) (gift from Yuichiro Miyaoka; Addgene; plasmid no. 108292; (Kato-Inui et al., 2018)).

#### Cell culture

A parental IMCD3-FlpIn cell line (gift from Peter K. Jackson, Stanford University, Stanford, CA) was modified to generate all stable cell lines used in the study. IMCD3-FlpIn cells were cultured in DMEM/F12 (11330-057; Gibco) supplemented with 10% FBS (100-106; Gemini Bio-products), 100 U/ml penicillin-streptomycin (400-109; Gemini Bio-products), and 2 mM L-glutamine (400-106; Gemini Bio-products).

The RPE1-hTERT cell line (ATCC CRL-4000) was cultured in DMEM/F12 supplemented with 10% FBS, 100 U/ml penicillin-streptomycin, 2 mM L-glutamine and 0.26% sodium bicarbonate (25080; Gibco).

The IMCD3 *Arl6*^−/−^ cell line was described previously (Liew et al., 2014; Nager et al., 2017). The genotype is NM_001347244.1:c.10_25del; c.3_6del.

Ciliation was induced by serum starvation in media containing 0.2% FBS for 16 to 24 h.

#### Transfection

For the generation of all stable cell lines, a plasmid encoding the Flp recombinase (pOG44) was co-transfected with the FRT-based plasmids using XtremeGene9 (Roche) via reverse transfection method into IMCD3 Flp-In cells as described (Liew et al., 2014). Stable transformants were selected by blasticidin resistance (4 μg/ml).

For CRISPR-based genome editing of TOM1L2, Cas9 and guide RNA were transiently expressed from a pX459 derivative and transfectants selected with puromycin. Clones were isolated by limited dilution and selected by western blotting. The genotype, NM_153080.3:c.174_175del;c.174_175insT, c.171_175del was determined by amplification of the targeted DNA region, DNA sequencing, and DECODR analysis (Deconvolution of Complex DNA Repair (Bloh et al., 2021)).

Transient transfection of pCilia-AMSH and pCilia-TOM1L2^BIR^ were performed using X-tremeGENE 9 DNA Transfection Reagent following the manufacturer’s protocol. 0.25μg of plasmid DNA was mixed with 0.75μl of transfection reagent (1:3 DNA: X-tremeGENE 9 ratio), incubated for 15 minutes at room temperature and then added onto 50,000 cells in suspension. The transfection-cell mixture was then transferred into one well of a 24-well plate.

For siRNA screens, 50,000 cells were transfected with indicated ON-TARGETplus siRNA SMARTpool duplexes using Lipofectamine RNAiMAX via reverse transfection. First, 1μl of RNAiMAX (13-778-030, Thermo Scientific) was diluted in 50 μl of Opti-MEM (31985070, Life Technologies) and incubated at room temperature for 5 min. Next, 0.5μl of RNAi duplex (20 pmol) were added to the diluted transfection reagent for 20 min before adding to the cells in suspension. The transfection-cell mixture was then transferred into one well of a 24-well plate. Cells were serum starved 48h later and processed for immunofluorescence another 16 h later. Cells were pretreated with 0.5μM CytoD for 10 min before the addition of sst (for SSTR3) for 6 h or SAG (for GPR161) for 3 h.

#### Antibodies and drugs

The following monoclonal antibodies were used for immunofluorescence: anti-acetylated tubulin (mouse; clone 6-11B-1; Sigma-Aldrich; 1:500), anti-ubiquitin clone FK2 (mouse; D058-3; Medical and Biological Laboratories; 1:500), anti-FLAG-M2 (mouse; F1804; Sigma-Aldrich), anti-Vps28 (mouse; sc-166537, clone E-7; Santa Cruz). The following polyclonal antibodies were used for immunofluorescence: anti-GPR161 (rabbit; 13398-1-AP; Proteintech; 1:100), anti-SMO (rabbit; a gift from Kathryn Anderson, Memorial Sloan Kettering Cancer Center, New York, NY; 1:500), anti-TOM1/TOM1L2 (rabbit; ab96320; Abcam; 1:500). The following monoclonal antibodies were used for immunoblotting: anti-Vps28 (mouse; sc-166537, clone E-7; Santa Cruz; 1:500), anti-FLAG-M2 (mouse; F1804; Sigma-Aldrich; 1:1000). The following polyclonal antibodies were used for immunoblotting: anti-MYO6 (rabbit; a gift from Folma Buss, Cambridge Institute for Medical Research, University of Cambridge, UK), anti-TOM1/TOM1L2 (rabbit; ab96320; Abcam; 1:500), anti-IFT122 (rabbit; ARP53817_P050; Aviva; 1:500), anti-IFT139 (rabbit; a gift from Pamela Tran, University of Kansas, USA), anti-IFT172 (rabbit; a gift from Kinga M. Bujakowska, Harvard Medical school, USA; 1:500), anti-IFT38 (rabbit; a gift from Hiroshi Hamada, RIKEN Center for Developmental Biology, Japan; 1:500), anti-BBS9 (rabbit; HPA021289: Sigma-Aldrich: 1:500), anti-BBS5 (rabbit; 14569-1-AP; Proteintech Group; 1:250), anti-BBS4 (rabbit; GN042; Maxence Nachury; 1:500), anti-LZTFL1 (rabbit; a gift from Val Sheffield, University of Iowa, USA; 1:500), anti-ARL6 (rabbit; Maxence Nachury; 1:500), anti-Actin (rabbit; A2066; Sigma-Aldrich: 1:1000), anti-TAX1BP1 (rabbit; HPA024432; Sigma-Aldrich: 1:1000), anti-HRS (rabbit; a gift from Harald Stenmark, University of Oslo, Norway; 1:500), anti-TOLLIP (rabbit; HPA038621; Sigma Aldrich; 1:500). Biotinylated SSTR3 and GPR161 were detected using Alexa Fluor 647-labeled monovalent streptavidin (mSA647) (Ye et al., 2018). The following reagents were used at the indicated concentrations: 200 nM SAG, 10 μM sst-14, and 0.5μM Cytochalasin D. Somatostatin 14 stocks were made in DMEM/F12 media, SAG and Cytochalasin D were dissolved in DMSO.

#### APEX labeling and proteomics

APEX labeling was performed as described previously (Mick et al., 2015). In brief, serum-starved cells were incubated in medium containing 0.5 mM biotin tyramide for 45 min before hydrogen peroxide was added to a final concentration of 1 mM. After 1 min, the medium was aspirated and cells were washed three times with 1x PBS supplemented with 10 mM sodium ascorbate, 10 mM sodium azide and 5 mM Trolox. Cells were lyzed in lysis buffer (6 M urea, 0.3 M NaCl, 1 mM EDTA, 10 mM sodium ascorbate, 10 mM sodium azide, 5 mM Trolox, 25 mM Tris/HCl pH 7.5) and equal protein concentrations of triplicate samples were diluted 10-fold in wash buffer (0.5% [v/v] Triton X-100, 0.1% [w/v] SDS, 0.3 M NaCl, 1 mM EDTA, 10 mM sodium ascorbate, 10 mM sodium azide, 5 mM Trolox, 25 mM Tris/HCl pH 7.5) and subjected to streptavidin chromatography. Biotinylated proteins were allowed to bind to streptavidin sepharose (Thermo Fisher) for 1 hr at room temperature, beads washed extensively with wash buffer and urea buffer (4 M urea, 10 mM Tris pH 7.5). Bound proteins were reduced and alkylated and eluted by on-bead Lys-C/trypsin digestion.

#### Proteomic profiling

*Arl6*^−/−^ and WT cilia proteins were compared after APEX labeling in triplicate. Isolated tryptic peptides were desalted by C18 SPE (Empore, 3M), dried under vacuum, and resuspended in 5% formic acid and 5% acetonitrile for analysis by LC/MS-MS on an Orbitrap Fusion mass spectrometer (Thermo Fisher Scientific) coupled to a Proxeon EASY-nLC II liquid chromatography (LC) pump (Thermo Fisher Scientific). Peptides were separated on a 75μm (inner diameter) microcapillary column packed with ~0.5 cm of Magic C4 resin (5μm, 100 Å, Michrom Bioresources) followed by ~35 cm of GP-18 resin (1.8 μm, 200 Å, Sepax) using a 1 hr gradient of 5-21% acetonitrile in 0.125% formic acid at a flow rate of ~500 nL/min. MS1 scans were detected in the Orbitrap with a resolution of 120,000, scan range of 400-1400 m/z, and maximum injection time of 100 ms. The most intense species from each MS1 was isolated in the quadrupole (isolation window 0.7) and fragmented by CID (collision energy 30%). MS2 spectra were detected in the Ion Trap using Ion Trap Rapid Scan Rate, a maximum injection time of 150 ms, and a normalized collision energy of 35.

Mass spectra were processed and converted to mzXML using a modified version of ReAdW.exe and searched against all entries from the mouse Uniprot database and the cilia-APEX fusion protein concatenated to a reverse sequence database using the SEQUEST algorithm using a 50 ppm precursor ion tolerance, trypsin protease specificity, and allowing for two missed cleavages. Peptides were filtered to a 1% FDR, and assembled proteins further filtered to a 1% FDR.

Raw spectral counts were used to calculate a normalized spectral abundance factor (NSAF) for each protein as previously described (Zybailov et al., 2006). Based on the distribution of log(NSAF) values, spectral counts of 0 were replaced by 0.25. The *Arl6*^−/−^ to WT ratios for each protein were determined by mean spectral counts calculated for each genotype. Proteins with mean spectral counts below 2.5 were excluded from analysis.

#### Imaging and microscopy

For fixed imaging, 50,000 cells were seeded on acid-washed 12 mm diameter #1.5 coverslips (12-545-81; Thermo Fisher Scientific), grown for 24 h, and then serum starved for 16 to 24 h before experimental treatment. After treatment, cells were fixed with 4% paraformaldehyde (50-980-487, Thermo Fisher Scientific) in PBS for 15 min at 37°C and extracted in −20°C methanol for 5 min (except for **Fig 2B and E**, where the methanol step was omitted). Cells were then permeabilized in IF buffer [PBS supplemented with 0.1% Triton X-100, 5% normal donkey serum (017-000-121; Jackson ImmunoResearch Laboratories), and 3% bovine serum albumin (BP1605-100; Thermo Fisher Scientific)]. Cells were then incubated at room temperature for 1h with primary antibodies diluted in IF buffer, washed three times in IF buffer, and then incubated with secondary antibodies (Jackson ImmunoResearch Laboratories) diluted in IF buffer for 30 min. Cells were then washed three times with IF buffer and DNA was stained with Hoechst 33258 (H1398; Molecular Probes). Cells were washed twice more with PBS, and coverslips were mounted on slides using fluoromount-G (17984-25; Electron Microscopy Sciences).

Cells were imaged either on a DeltaVision system (Applied Precision) equipped with a PlanApo 60x/1.40 objective lens (Olympus), CoolSNAP HQ2 camera (Photometrics), and solid-state illumination module (InsightSSI) or on confocal LSM 700 or LSM 900 (Zeiss) microscopes equipped with 40x Plan-Apochromat 1.3 DIC oil objective. Z stacks were acquired at 0.2 μm interval (DeltaVision) except for **Fig. 2B** (LSM 700), **Fig. 7C,E** (LSM900), (0.5 μm interval). Except for **Fig. 2B** (LSM700) and **Fig. 8 A and C** (LSM900), all images were acquired using the DeltaVision microscope. Line scans of ^NG3^BBS5 fluorescence along cilia were generated by capturing images of cilia via total internal reflection microscopy (TIRF). TIRF illumination reduced background fluorescence and increased the signal to noise ratio of ciliary signals to background signals. Fixed cells were imaged using a Plan Apochromat 60× 1.49 NA TIRF oil objective lens (Olympus) and a 488-nm laser from DeltaVision Quantifiable Laser Module (50% laser power). Z stacks were acquired at 0.2 μm interval and most in-focus planes were used for representative images in **Fig. 3A-B, Fig. 3C-D, and Fig. 5 D-E.**

#### Image analysis

##### Measurement of ciliary signals and of GPCR exit index

Files were imported from the Deltavision or LSM700/900 workstations into ImageJ/ Fiji (National Institutes of Health) for analysis. For the quantification of ciliary signals (for all the proteins, in all the figures) in fixed cells, maximum intensity projections were used. The ciliary intensities were measured using the following equation:

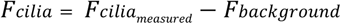

*Fcilia_measured_* is the total ciliary fluorescence detected, *Fbackground* is the background fluorescence measured in the adjacent area. Ciliary fluorescence was measured in ImageJ by a plot profile of a 3-pixel-wide line along the long axis of the cilium and the same line was used to measure the fluorescence in the adjacent area. For all measurements, the fluorescence integrated density was used.

To generate the heatmap representing the ciliary exit of GPCRs in RNAi screens, the following formula was used:

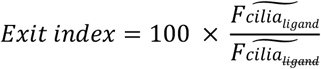

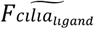 is the median of the *Fcilia_medsured_* upon ligand treatment (+sst for SSTR3/ +SAG for GPR161), 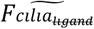 is the median of the *Fcilia_measured_* in untreated conditions (-sst for SSTR3/ -SAG for GPR161).

No gamma adjustment was applied during figure preparation. For some representative micrographs, the most in-focus plane was used rather than the maximum intensity projection (Fig.2E, Fig.5B and F, Fig.6C, Fig.7 C and E).

Integrated fluorescence intensities were used for all the measurements. The ciliary intensities (*Fcilia*) were plotted as violin plots using the PlotsOfData web tool (https://huygens.science.uva.nl/PlotsOfData/) (Postma and Goedhart, 2019). Each violin represents the distribution of data, including all the data points. Median and interquartile range are marked by solid and dotted lines, respectively.

###### Linescans

Linescan were generated as described (Ye et al., 2018). Line scans were generated by measuring longitudinal fluorescence intensities of BBS5 in ImageJ by a plot profile of a 5-pixel-wide line along the long axis of cilia. Data from multiple cilia were averaged by assigning a length percentage to the pixel intensities. 0% referred to the base, and 100% referred to the tip; resulting values were then grouped into 5% bins and averaged. Bin means from multiple cilia were averaged and plotted.

##### *Chlamydomonas* culture and immunofluorescence

*Chlamydomonas* WT CC-4533 cw15 mt-Jonikas CMJ030], and *Tom1* (*Cre06.g292000*) LMJ.RY0402.161500, and, LMJ.RY0402.134895, and, *Bbs4*-CC-4377 ptx5-1/bbs4-1:: NIT1 agg1 mt+ (*Chlamydomonas* Resource Center) strains were cultured under 14:10 h light and dark cycle in TAP media (Gorman and Levine, 1965) for 72 h.

*Chlamydomonas* immunofluorescence was performed as follows. Cells were fixed with 4 % paraformaldehyde (50-980-487, Thermo Fisher Scientific) in MT buffer (30 mM HEPES, pH7.0, 5 mM EGTA, 5 mM MgSO_4_, and 25mM KCl) for 20 minutes in suspension. The cells were centrifuged in a Beckman SX4750A rotor at 1000 rpm for 5 minutes, resuspended in ~100 μl of fixative, and transferred onto slides coated with 0.1% polyethyleneimine (PEI, 9002-98-6, 26913-06-4, Polyethylenimine, Linear, MW 25000). After 2-3 minutes, the unadhered cells were washed off by rinsing with PBS. Cells were permeabilized in 0.5% Triton-X 100 for 20 minutes followed by blocking for 1 h in blocking buffer (3% fish skin gelatin, 1% BSA, 0.1% Tween-20 in PBS) at room temperature. Cells were incubated at 4°C overnight with primary antibodies diluted in blocking buffer, washed five times in PBS, and incubated with secondary antibodies for 2 h at room temperature. After five washes in PBS, cover glasses were mounted on slides using Fluoromount-G. Cells were imaged on the LSM900 confocal microscope.

##### Recombinant protein expression and GST capture assays

GST-tagged TOM1, TOM1L1, TOM1L2 FL and truncations, TOM1L2 mutants, Myo6-CBD, and mNedd4△N52 protein fusions were expressed in Rosetta2(DE3)-pLysS cells grown in 2xYT medium (Millipore Sigma, Y2627) at 37°C until the optical density (OD) at 600 nm reached 0.6. Protein expression was then induced with 0.2 mM IPTG at 18°C for 16 h. Post-induction the cells were pelleted down at 6,000 x *g* for 15 min at 4°C and the pellets were resuspended in 4XT (200 mM Tris, pH 8.0, 800 mM NaCl, 1 mM DTT) supplemented with protease inhibitors (1 mM AEBSF, 0.8 μM Aprotinin, 15 μM E-64, 10 μg/mL Bestatin, 10 μg/mL Pepstatin A and 10 μg/mL Leupeptin) and lysed by sonication. The lysates were clarified by centrifugation at 30,000 x *g* for 30 min at 4°C. The clarified lysates were loaded onto Glutathione Sepharose 4B resin (Cytiva) and proteins eluted with 50 mM reduced glutathione in buffer XT (50 mM Tris, pH 8.0, 200 mM NaCl, 1 mM DTT). In some cases, the GST tag was cleaved off the fusion protein by incubating the protein bound to the glutathione resin with Prescission Protease (0.5μg/μl, Cytiva) in one-bed volume of 2XT buffer. Proteins were subsequently dialyzed against XT buffer with one change of buffer and flash-frozen in liquid nitrogen after the addition of 5% (w/v) glycerol.

N-terminally GST-tagged ARL6ΔN16[Q73L] was expressed in bacteria as described (Jin et al., 2010). The BBSome was purified from bovine retina by ARL6^GTP^-affinity chromatography as described (Chou et al., 2019) and was used in the GST capture assays immediately after purification.

For mapping of the BBSome-binding region on TOM1L2, GST capture assays were performed as follows: 100 μg of purified GST-TOM1L2 (or truncations or point mutants) were immobilized onto 10 μl Glutathione Sepharose 4B resin and were incubated with purified BBSome (0.7 to 1 μg) at 4°C for 1h in LepRb buffer (20 mM HEPES-NaOH, pH 7.0, 5 mM EDTA, 20 μg/ml glycerol, 300 mM KOAc, 1 mM DTT, 0.2% Triton X-100) supplemented with protease inhibitors. The beads were washed in LepRb buffer and the bound proteins were eluted by boiling beads in 30 μl LDS loading buffer and loaded onto the SDS-PAGE gels for western blot analysis.

For mapping of the MYO6 binding region on TOM1L2, GST capture assays were performed as follows: 100 μg of purified GST-TOM1L2 (or truncations or point mutants) were bound to 10 μl Glutathione Sepharose 4B resin and were incubated with purified MYO6-CBD (100 μg) at 4°C for 1h in NSC250 buffer (25mM Tris pH 8, 250mM KCl, 5mM MgCl2, 1mM DTT, 0.5% CHAPS) supplemented with protease inhibitors. The beads were washed in NSC250 buffer and subjected to cleavage elution by PreScission protease (7.5 μg). Eluates were boiled in LDS loading buffer before loading onto SDS-PAGE gels for Coomassie analysis.

##### Synthesis of UbK63 chains and binding assays

The autoubiquitination of NEDD4 was performed as described in (Kim et al., 2007). Briefly, 100μg of GST-NEDD4△52 was bound to 10 μl Glutathione sepharose resin and incubated with 50 nM ubiquitin activating enzyme UBA1 (R&D Systems, E305025), 750 nM ubiquitin conjugating enzyme UBCH5B (R&D Systems, E2622100), 2 mM ATP, and 58 μM ubiquitin (R&D Systems, U-100H-10M) in 50 μl reaction buffer (20 mM Tris-Cl pH 7.6, 20 mM KCl, 5 mM MgCl_2_, and 1mM DTT) at 37°C for 60 min. Beads were washed three times in NSC250 buffer to remove the unbound enzymes and ubiquitin. The binding of proteins to UbK63 chains grown onto GST-NEDD4 was performed by incubating the immobilized chains with either retinal extracts (**Fig. 1 B-C**), or IMCD3 cell extracts (**Fig. 1E**, **Fig. S1A**) or pure proteins (**Fig. 1D, Fig. S1B**). Beads were washed thrice in NSC250 buffer and bound material was eluted by cleaving ubiquitin linkages with 250 nM USP2cc in 30 μl of NSC250 buffer at 37°C for 90 min. Eluates were collected and boiled in LDS sample buffer before loading onto SDS-PAGE gels for western blot analysis.

##### Visual capture assays

HEK293 cells were transfected with and equimolar mixture of pCS2-GFP-BBS1, -BBS2, -BBS4, -BBS5, -BBS7, -BBS8, -BBS9, and - BBS18. 24h post-transfection cells, were trypsinized and pelleted down. Cell pellets were resuspended in ice-cold lysis buffer (20mM HEPES-KOH pH 7.4, 150 mM NaCl, 0.1% Triton X-100, 10% glycerol, 5mM DTT) supplemented with protease inhibitors (1 mM AEBSF, 0.8 mM Aprotinin, 15 mM E-64, 10 mg/mL Bestatin, 10 mg/mL Pepstatin A and 10 mg/mL Leu-peptin), incubated on ice for 15 min and spun down at 16,000 x *g* for 15 min at 4°C. Supernatants were collected and incubated with GST-TOM1L2 (or truncations or point mutants) bound to glutathione sepharose resin at 4°C for 1h. The beads were washed four times in lysis buffer. After the last wash, the beads were resuspended in 50μl of lysis buffer and 10μl of the resulting slurry were spotted onto a slide and mounted under a coverslip. Beads were imaged on a Zeiss Axiophot microscope equipped with InSight Spot camera (Diagnostic Instruments) and a 20X objective.

