## Supplemental Figures for "The ancestral ESCRT protein TOM1L2 selects ubiquitinated cargoes for retrieval from cilia"

**AFFILIATION:**

**SUPPLEMENTAL INFORMATION**

Figures S1-S9

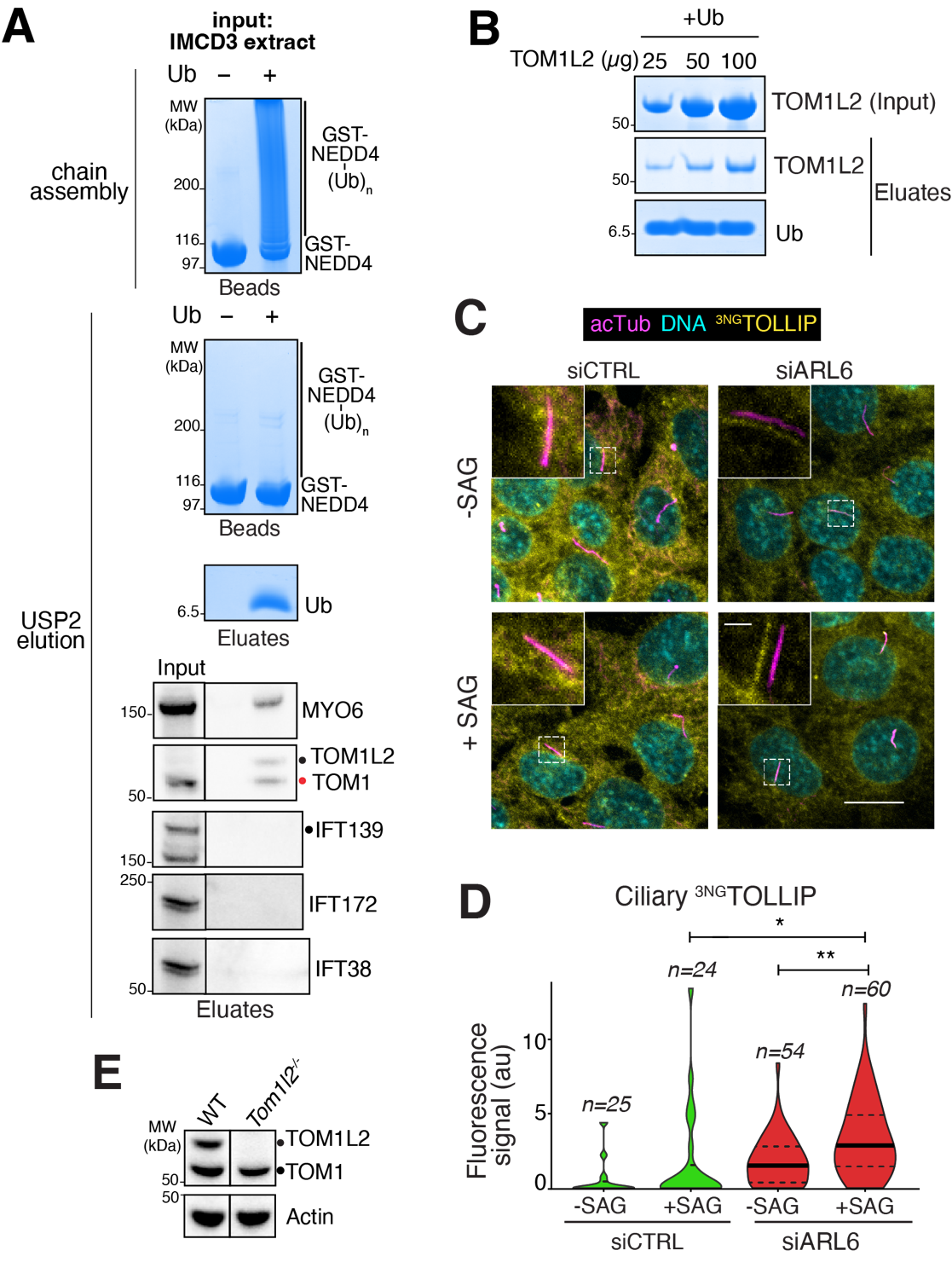

**Figure S1. UbK63 binding assays and TOLLIP immunofluorescence.**

**A.** Capture of UbK63 chain readers from IMCD3 cell extracts. Top panel: Coomassie-stained gel of bead-bound GST-NEDD4 after completion of autoubiquitination reaction. Ub was omitted from the ubiquitination reaction for the control. Middle and lower panels show the products of the treatment with USP2 after IMCD3 cells extracts were applied onto GST-NEDD4 ± (Ub)_n_ beads. Middle panel: Coomassie-stained gel of bead-bound GST-NEDD4 ± (Ub)_n_. Lower panel: Coomassie-stained gel of material released from beads by USP2 treatment. Bottom panels: western blots of known UBP and ciliary trafficking components. Efficient binding is detected to the known UbK63 readers Myosin VI (MYO6) and TOM1/TOM1L2. No binding to UbK63 chains is detected for the IFT-A complex subunit IFT139 or the IFT-B complex subunits IFT172 and IFT38. 0.1% input and 42% of cleavage eluates were loaded for the western analyses. **B.** Direct binding of TOM1L2 to UbK63 chains. Different concentrations of bacterially expressed TOM1L2 were incubated with immobilized UbK63 chains and bound material was eluted via USP2 mediated cleavage before SDS-PAGE Coomassie staining. 0.1% input and 42% capture eluates were loaded in each lane. **C.** IMCD3 cells stably expressing TOLLIP fused to triple tandem mNeonGreen (3NG) were transfected with siRNAs targeting ARL6 (siARL6) or negative control siRNA (siCTRL), and cells were treated with or without Smoothened Agonist (SAG) for 2 h, then fixed and stained for acetylated tubulin (acTub, magenta). ^3NG^TOLLIP was imaged through the intrinsic fluorescence of NG (yellow). Scale bars: 5μm (main panel), 1μm (inset). A weak ciliary TOLLIP signal detected in WT cells is markedly elevated in ARL6-depleted cells. **D.** The ciliary fluorescence intensity of the TOLLIP channel was measured in each condition, and the data are represented as violin plots. Asterisks indicate statistical significance values calculated by one-way ANOVA followed by Tukey’s post hoc test. **, *p* ≤ 0.01; *, *p* ≤ 0.05. *n* = 24-60 cilia. **E.** Protein extracts from WT and *Tom1l2^-/-^* cells were resolved on SDS-PAGE and probed for TOM1/TOM1L2 and actin.

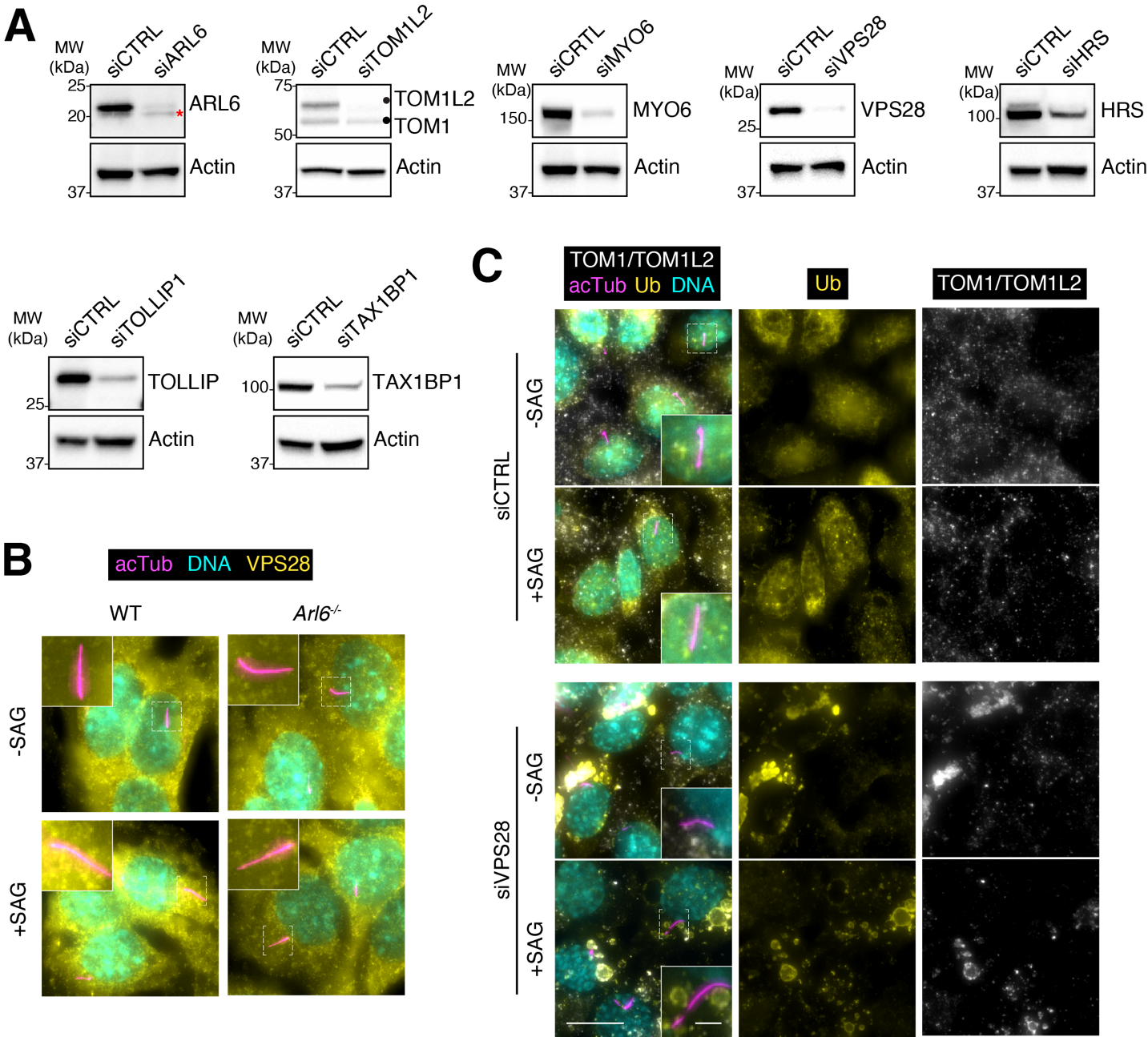

**Figure S2. Validation of siRNA-mediated protein depletion and characterization of VPS28-depleted cells.**

**A.** Lysates from cells treated with the indicated siRNA used in the GPCR exit screens were immunoblotted for the target proteins and for actin as a loading control. A non-specific band cross-reacting with the anti-ARL6 antibody is marked with an asterisk. A commercial antibody raised against TOM1 detects both TOM1 and TOM1L2 (see **Tumbarello, D. A. et al.** Nature Cell Biology 14:1024–1035 (2012) for additional validation of this antibody). **B.** WT or *Arl6 ^-/-^* IMCD3 cells were treated with or without SAG for 2 h, fixed, and stained for acetylated tubulin (acTub, magenta) and VPS28 (yellow). Scale bar: 5μm (main panel), 1μm (inset). VPS28 is detected on endomembrane structures but no VPS28 signal is detected inside cilia. **C.** IMCD3 cells were transfected with control or VPS28 siRNA, treated with or without SAG for 2 h, fixed, and stained for acetylated tubulin (acTub, magenta), TOM1/TOM1L2 (white), and ubiquitin (yellow). Scale bar: 5μm (main panel), 1μm (inset). VPS28 depletion results in the accumulation of TOM1/TOM1L2 and ubiquitin on the endosomal structures.

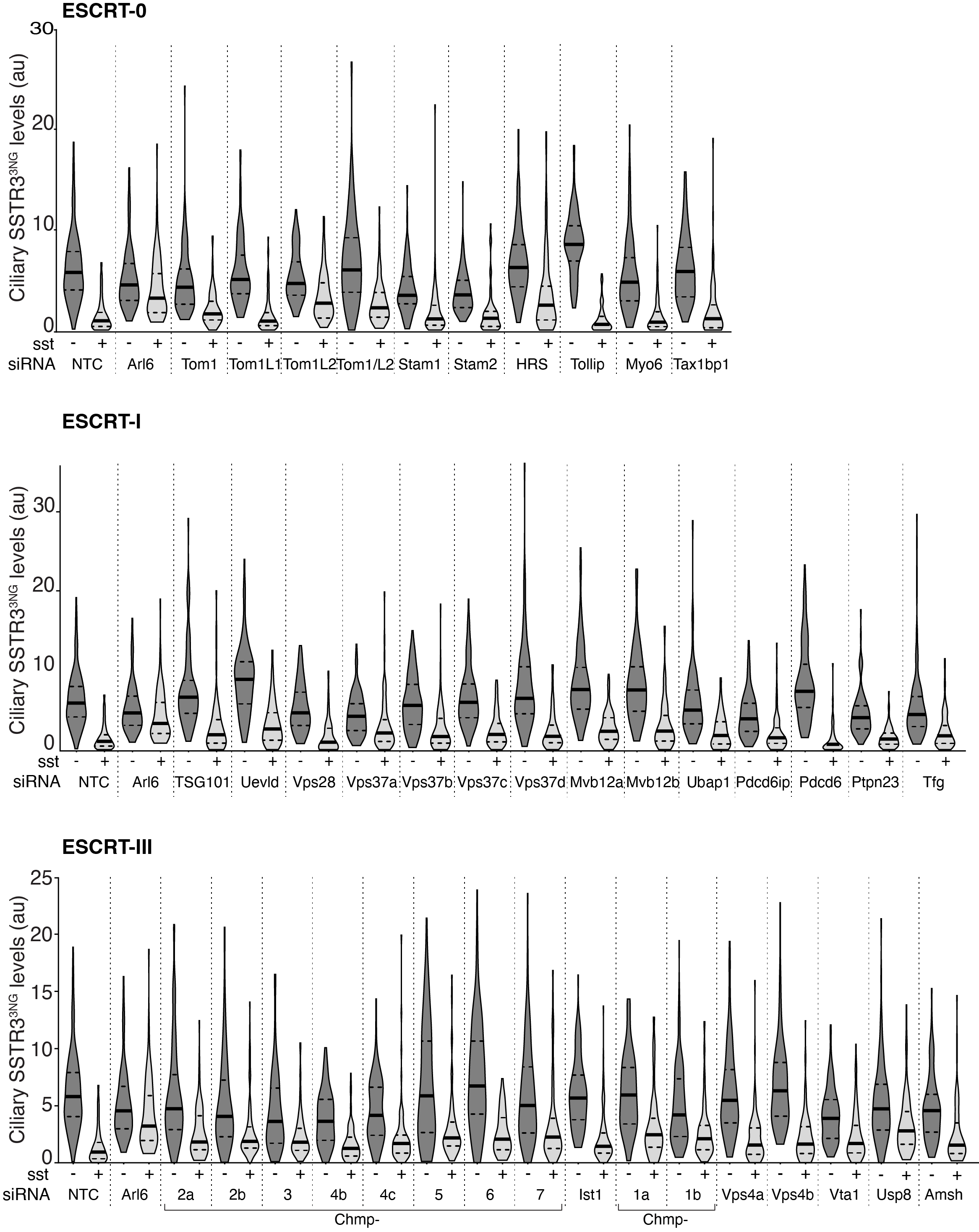

**Figure S3. siRNA screen of ESCRT and candidate UbK63 readers for SSTR3 exit from cilia.**

IMCD3-^AP^SSTR3^NG^ cells were transfected with indicated siRNAs, treated with or without sst for 6h in presence of 0.5 µM Cytochalasin D, fixed and stained for acetylated tubulin. ^AP^SSTR3^NG^ was imaged through the intrinsic fluorescence of NG. The fluorescence intensity of ^AP^SSTR3^NG^ was measured in each condition, and the data are represented as violin plots. *n* = 80-120 cilia.

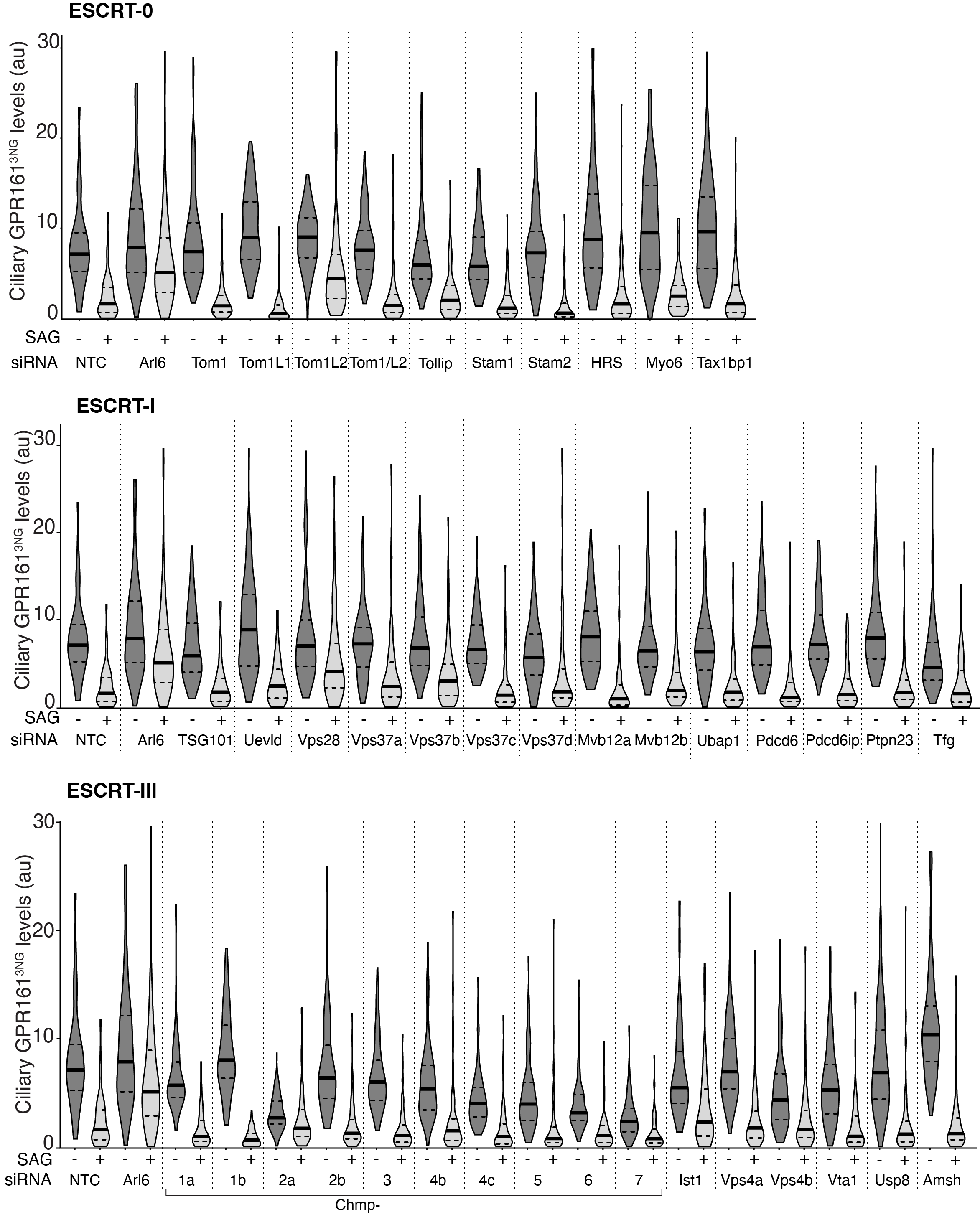

**Figure S4. siRNA screen of ESCRT and candidate UbK63 readers for GPR161 exit from cilia.**

IMCD3-^AP^GPR161^NG3^ cells were transfected with indicated siRNAs, treated with or without SAG for 3h in presence of 0.5µM Cytochalasin D, fixed, stained for acetylated tubulin. ^AP^GPR161^NG3^ was imaged through the intrinsic fluorescence of NG. The fluorescence intensity of ^AP^GPR161^NG3^ was measured in each condition, and the data are represented as violin plots. *n* = 80-120 cilia.

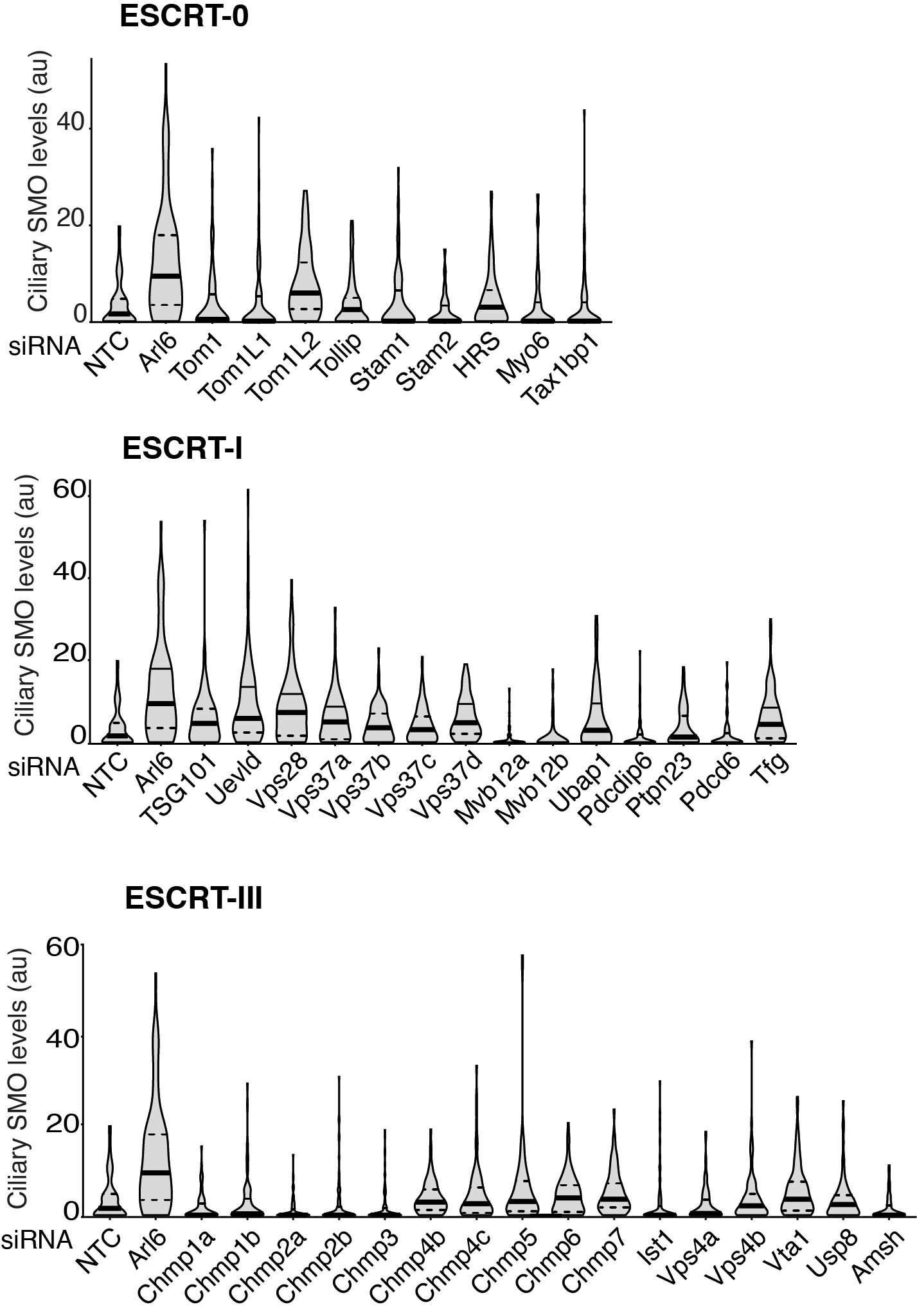

**Figure S5. siRNA screen of ESCRT and candidate UbK63 readers for SMO exit from cilia.**

IMCD3- ^AP^GPR161^NG3^ cells were transfected with indicated siRNAs, treated with or without SAG for 3h in presence of 0.5µM Cytochalasin D, fixed, stained for acetylated tubulin and SMO. ^AP^GPR161^NG3^ was imaged through the intrinsic fluorescence of NG. The fluorescence intensity of SMO was measured in each condition, and the data are represented as violin plots. *n* = 80-120 cilia.

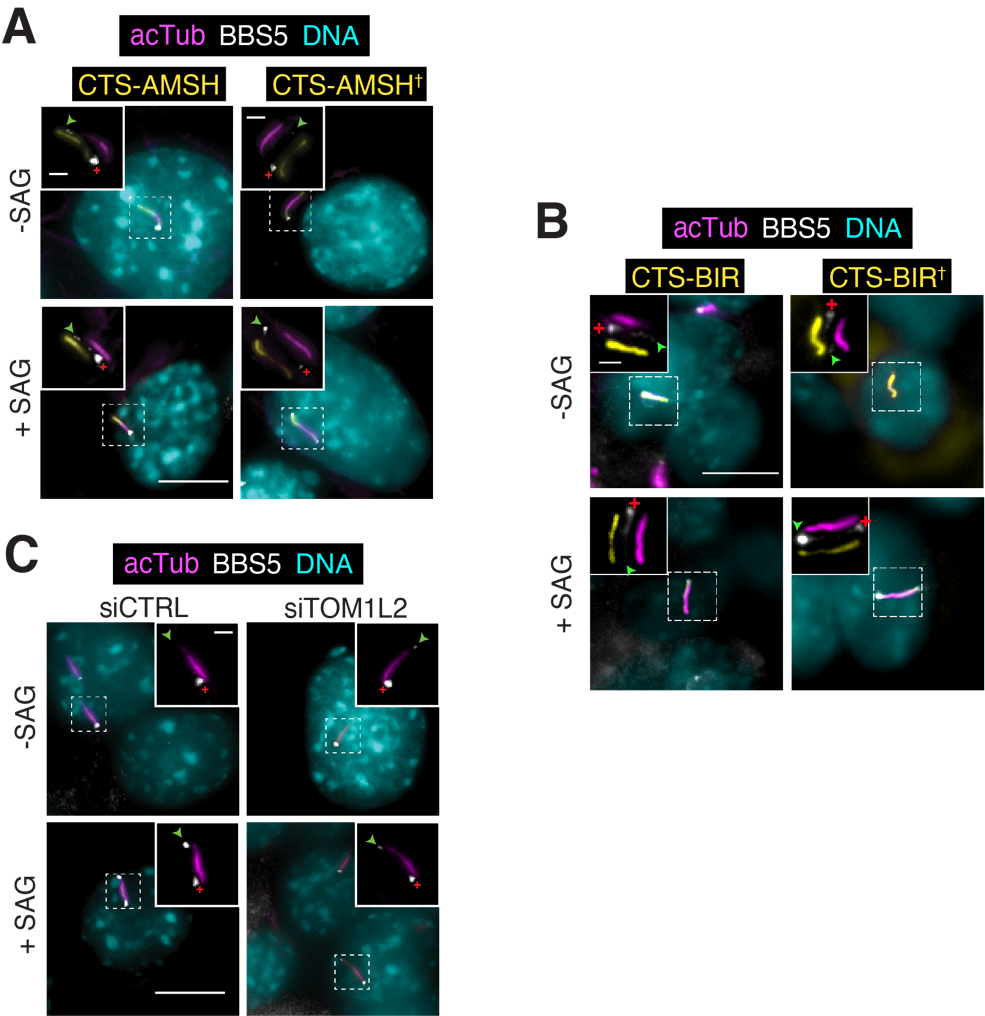

**Figure S6. BBSome staining in cells expressing retrieval disrupting agents.**

**A-B.** IMCD3-[pEF1α-^NG3^BBS5] were transfected with plasmids expressing either AMSH (**A**) or the BBSome interacting region of TOM1L2 (BIR, **B**) or indicated variants fused to the ciliary targeting signal of NPHP3 (CTS) and mScarlet. Cells were then treated with or without SAG for 40 min, fixed and stained for acetylated tubulin (acTub; magenta), and DNA (cyan). ^3NG^BBS5 was imaged through intrinsic fluorescence of NG and CTS through mScarlet. Representative images of cilia are shown in insets. Scale bar: 5μm (main panel), 1μm (inset). White crosses mark the location of the basal body, and an arrowhead marks the tip of the cilia. **C.** IMCD3-[pEF1α-^NG3^BBS5] were transfected with control or TOM1L2 siRNA, treated with or without SAG for 40 min, fixed, and stained for acetylated tubulin (acTub, magenta), and DNA (cyan). Scale bar: 5μm (main panel), 1μm (inset).

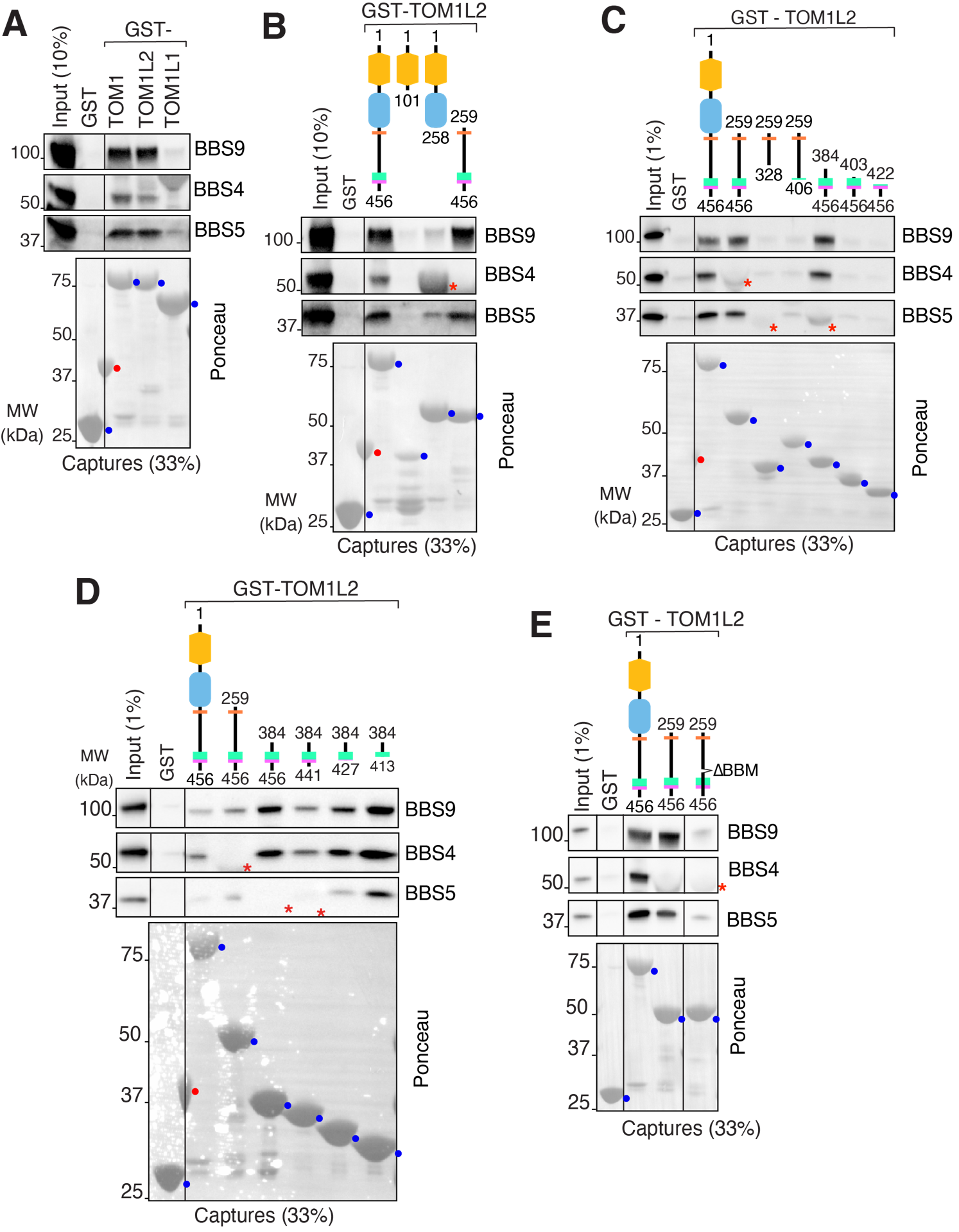

**Figure S7. Mapping of the BBSome-binding determinant on TOM1L2.**

**A.** The capture assays shown in **Fig. 4B** were immunoblotted for BBS4 and BBS5 and total proteins stained with Ponceau S. The BBS9 immunoblot shown in **Fig. 4B** is included. **B.** Additional immunoblots and Ponceau stain for **Fig. 4C**. **C.** Additional immunoblots and Ponceau stain for **Fig. 4D**. **D.** Additional immunoblots and Ponceau stain for **Fig. 4E**. **E.** Deletion of the BBM in TOM1L2 leads to a drastic reduction of BBSome binding. In all Ponceau stain panels, the GST-TOM1 fusions proteins are marked with a blue dot and a red dot marks a protein from neighboring lanes. In some panels, an asterisk indicates GST-tagged proteins detected by immunoblot that interfere with detection of a BBSome subunit.

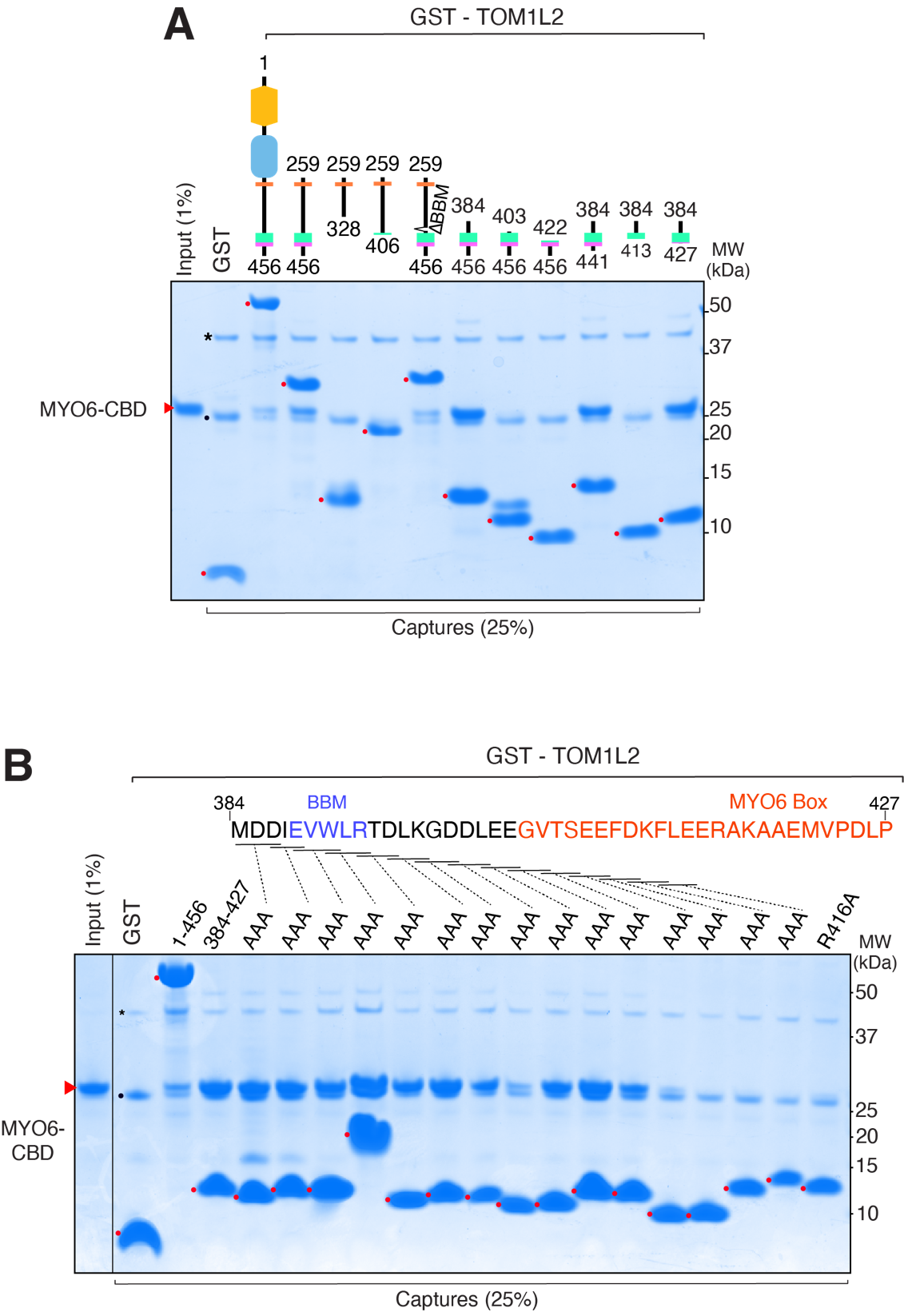

**Figure S8. Mapping of the MYO6-binding determinant on TOM1L2.**

**A.** Deletion mapping of the MYO6-binding region of TOM1L2. The cargo binding domain (CBD; aa 1060-1253) of Myosin VI was expressed in bacteria and purified to near-homogeneity. GST-TOM1L2 truncations were immobilized on glutathione Sepharose and beads were incubated with MYO6-CBD. Bound material was eluted via PreScission cleavage and eluates were resolved on SDS-PAGE and stained with Coomassie. The cleaved TOM1L2 fragments are marked with red dots and the red arrowhead marks MYO6-CBD. PreScission is marked by an asterisk, and a black dot indicates the position of GST which contaminates the eluates. 1% input and 25% of eluates were loaded for Coomassie analysis. The weak binding of MYO6 to the truncations encompassing the complete CTD of TOM1L2 is greatly increased with the shorter truncations that start at position 384, likely indicating autoinhibition of MYO6 binding within the CTD of TOM1L2. The minimal MYO6-binding region of TOM1L2 spans aa 384-427. **B.** Mapping of the MYO6-binding determinant of TOM1L2 by alanine scanning mutagenesis. Capture assays of the MYO6-CBD were conducted with triple alanine mutants of TOM1L2^384-427^ covering aa 384-414. The BBM defined in **Fig. 4** is in blue and the previously defined MYO6-binding helix (**Hu et al.** Nat Comm. 10: 3459 (2019)) is in red. Our mapping is in close agreement with the study by Hu et al. with the additional finding that ^398^DD^399^ may provide an additional binding surface to MYO6.

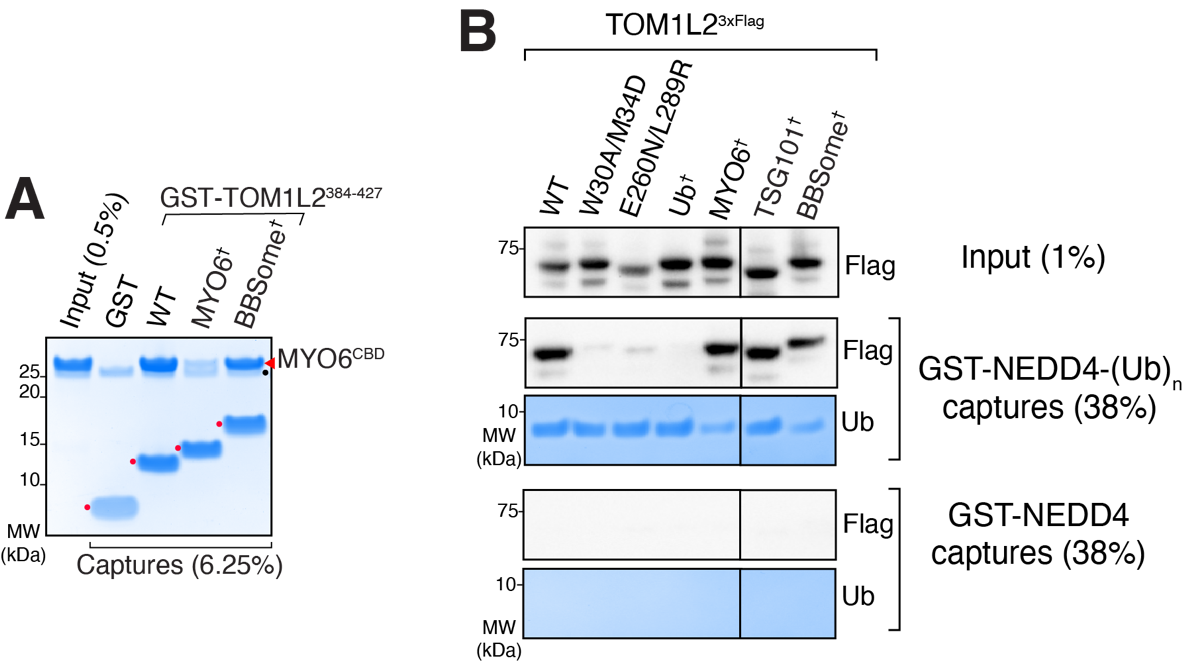

**Figure S9. Validation of TOM1L2 variants**

**A.** Capture assays of the MYO6 cargo binding domain (CBD) by GST-TOM1L2 variants confirm that the L464E/R467E mutation greatly reduces TOM1L2 binding to MYO6. Assays were conducted as in **Fig. S8**. The cleaved TOM1L2 fragments are marked with red dots, the red arrowhead marks MYO6-CBD, and a black dot indicates the position of GST protein that contaminates the eluates. **B.** Capture assays with UbK63 chains grown onto GST-NEDD4 find greatly diminished UbK63 binding by mutations in the VHS or GAT domain and undetectable UbK63 chain binding when both mutations in both domains are combined. Assays were conducted as in **Fig. 1E**. GST-NEDD4 without Ub in the chain-building reaction served as a control.
